# CNS-derived SerpinA3n restricts pathogenesis during WNV encephalitis via suppression of inflammatory protease activity

**DOI:** 10.64898/2026.11.26.769430

**Authors:** Vishal Karuppusamy, Eduard Marmut, Juan P. Angel, Evan DaPrano, Colm Atkins, Brian P. Daniels

## Abstract

Proteolytic activity is integral to host immune defense, yet little is known about how proteolytic homeostasis is maintained in the central nervous system (CNS) during viral infection. Here, we investigated roles for the serine protease inhibitor SerpinA3n in promoting host protection during West Nile virus (WNV) encephalitis. WNV neuroinvasion was accompanied by widespread induction of serpin expression, including SerpinA3n. Pharmacological inhibition of SerpinA3n within the CNS, but not in the periphery, exacerbated mortality, whereas CNS supplementation with recombinant SerpinA3n improved survival, preserved blood-brain barrier (BBB) integrity, and reduced proteolytic activity and leukocyte accumulation without altering CNS viral burden. Using cell-type-specific genetic deletion, we identified astrocytes and neurons as major sources of SerpinA3n in the infected CNS. Loss of SerpinA3n from either cell population increased mortality, BBB permeability, and gelatinase/collagenase activity. However, loss of neuronal, but not astrocytic, SerpinA3n exacerbated leukocyte accumulation and antigen-presenting cell activation, suggesting that the cellular context of SerpinA3n production influences its immunoregulatory functions. Both astrocyte- and neuron-derived SerpinA3n restrained granzyme B activity in the infected CNS, and neutralization of granzyme B reversed the excess mortality associated with pharmacological inhibition of SerpinA3n. Together, these findings identify serpin-mediated regulation of inflammatory protease activity as a mechanism of host protection during viral encephalitis and establish both neurons and astrocytes as important contributors to the maintenance of proteolytic homeostasis in the inflamed CNS.

## Introduction

Host immune responses to viral infections of the central nervous system (CNS) must balance the need to restrict pathogen replication with the preservation of neural tissue. Although inflammatory signaling pathways and the contributions of individual immune cell populations to viral neuropathogenesis have been extensively studied, comparatively little is known about the mechanisms that maintain protein homeostasis (proteostasis) in the CNS during infection. In particular, the regulation of proteolysis may be a critical determinant of tissue integrity in the inflamed CNS, where the release of proteolytic enzymes by resident CNS and infiltrating immune cells threatens the stability of extracellular matrix, neurovascular, and neuronal proteins [1–3]. Defining the endogenous mechanisms that regulate protease activity during CNS infection may therefore reveal important determinants of host susceptibility to virus-induced neurologic disease.

Proteolysis is central to the initiation, amplification, and resolution of immune responses [4,5]. Serine proteases, in particular, participate in diverse immunologic processes, including complement activation, coagulation, cytokine regulation, and cell-mediated cytotoxicity [6,7]. In addition to directly cleaving structural and signaling proteins, serine proteases can initiate or amplify proteolytic cascades through the activation of other proteases synthesized as inactive zymogens, including matrix metalloproteinases and caspases [8–10]. Thus, the consequences of dysregulated serine protease activity may extend well beyond the cleavage of their immediate substrates. In the CNS, where the maintenance of specialized intercellular junctions, extracellular matrix architecture, and neuronal connectivity is essential for normal tissue function [11], disruption of proteolytic homeostasis may have particularly deleterious consequences.

Among the serine proteases most prominently associated with inflammatory tissue injury are granzyme B, neutrophil elastase, and cathepsin G, which are released by activated cytotoxic lymphocytes and neutrophils [12,13]. Although granzyme B is best characterized for its role in the induction of apoptotic cell death [10], it also acts extracellularly to cleave basement membrane and junctional proteins and to modulate neuronal signaling through protease-activated receptors [14–17]. Granzyme B has been implicated in neuronal injury and neurodegeneration during CNS infection with tick-borne encephalitis virus and cerebral malaria [18,19]. Similarly, neutrophil elastase and cathepsin G contribute to blood-brain barrier (BBB) dysfunction and tissue injury in models of CNS autoimmunity, cerebral ischemia, and traumatic brain injury [20–23]. These observations suggest that proteases mobilized as components of the host immune response may be important mediators of neuropathology across diverse inflammatory insults to the CNS.

Endogenous regulation of protease activity is mediated in part by serine protease inhibitors (serpins), a diverse superfamily of proteins that regulate proteolysis across numerous physiological and immunological processes [24]. Inhibitory serpins function as suicide substrates, engaging their target proteases through a reactive center loop and undergoing a conformational rearrangement that results in irreversible inhibition of the protease [25]. By restraining the activity of individual proteases, serpins can also limit the propagation of downstream proteolytic cascades. Although several serpins have been implicated in the pathogenesis of CNS disorders characterized by neuroinflammation, including ischemic stroke, traumatic brain injury, and Alzheimer’s disease [26], the contributions of this protein family to the maintenance of proteostasis during CNS infection remain poorly understood. Moreover, the cellular sources of serpins within the CNS and the extent to which distinct neural cell populations contribute to the local regulation of inflammatory proteases are largely undefined.

SerpinA3n, the murine homolog of human SERPINA3, inhibits several inflammation-associated serine proteases, including granzyme B, neutrophil elastase, and cathepsin G [27–29]. SerpinA3n has been ascribed both protective and pathogenic functions in models of sterile neuroinflammation, suggesting that its biological effects may depend on its cellular source and the inflammatory context in which it is expressed [30]. We previously identified SerpinA3n as a component of a RIPK3-dependent transcriptional program induced in astrocytes during Zika virus (ZIKV) encephalitis [31]. Supplementation of SerpinA3n in the CNS preserved BBB integrity and improved survival in infected mice, implicating the local regulation of proteolysis as a potentially important mechanism of host protection during viral encephalitis [31]. However, the extent to which endogenously expressed serpins regulate inflammatory protease activity in the infected CNS, and how the compartmentalization of serpin expression among neural cell populations influences disease pathogenesis, remain unresolved.

West Nile virus (WNV), a mosquito-borne member of the genus *Orthoflavivirus*, is a leading cause of arboviral encephalitis. The geographic distribution of WNV and other orthoflaviviruses is expanding in association with climate-driven changes in vector ecology [32,33], and there are currently no approved human vaccines or antiviral therapies for WNV infection [32]. Although neuroinvasive disease develops in fewer than 1% of infected individuals, WNV encephalitis is associated with substantial mortality and neurologic morbidity [34]. Following neuroinvasion, recognition of viral pathogen-associated molecular patterns (PAMPs) by innate immune sensors in infected neurons initiates inflammatory cytokine and chemokine responses [35–38], which are amplified by activated glia [36,39,40] and promote the recruitment of peripheral leukocytes [41]. Although these responses are essential for viral clearance [42], they can also contribute to tissue injury [43]. Granzyme B-expressing cytotoxic lymphocytes are important components of the antiviral immune response to WNV [44–47], yet the endogenous mechanisms that constrain the activity of granzyme B and other inflammatory proteases within the infected CNS remain poorly characterized.

Here, we investigated the contribution of serpin-mediated regulation of proteolysis to host protection during viral encephalitis using a mouse model of WNV infection. We found that CNS invasion by WNV was accompanied by the induction of several serpins, including SerpinA3n, and that pharmacological inhibition of SerpinA3n within the CNS, but not in the periphery, exacerbated mortality. Conversely, supplementation of SerpinA3n in the CNS improved survival, preserved BBB integrity, reduced gelatinase and collagenase activity, and attenuated leukocyte accumulation and activation without altering CNS viral burden. Using cell-type-specific genetic deletion, we identified both astrocytes and neurons as major sources of SerpinA3n in the infected CNS and demonstrated that loss of SerpinA3n from either cell population increased mortality, BBB permeability, and proteolytic activity in the CNS. Notably, loss of neuronal, but not astrocytic, SerpinA3n also exacerbated CNS leukocyte infiltration and activation, suggesting that the spatial and cellular context of SerpinA3n production may confer distinct immunoregulatory functions. Finally, we identified granzyme B as a critical target of endogenous SerpinA3n activity in the infected CNS and demonstrated that neutralization of granzyme B reversed the excess mortality associated with SerpinA3n inhibition. Collectively, these findings establish the local regulation of inflammatory protease activity as an important mechanism of host protection during viral encephalitis and identify both neurons and astrocytes as active contributors to the maintenance of proteolytic homeostasis in the inflamed CNS.

## Results

### Subcutaneous WNV infection in mixed C57BL/6J × 129S mice results in CNS dissemination and induction of SerpinA3n expression

To investigate the contribution of endogenous SerpinA3n to host protection during viral encephalitis, we examined the pathogenesis of WNV infection in mice with cell-type-specific deficiencies in *SerpinA3n*. The *Serpina3n*^fl/fl^ mouse line used in these studies was previously generated on a mixed C57BL/6J × 129S genetic background. Given the influence of host genetics on susceptibility to orthoflavivirus neuroinvasion [48,49], we first characterized the kinetics of WNV dissemination following subcutaneous infection of wild-type mice on this background **(Fig. 1A)**. WNV RNA levels were significantly elevated in the cortex at 7 and 9 days post-infection (dpi) **(Fig. 1B)** and in the brainstem, cerebellum, and spinal cord at 9 dpi **(Fig. 1C–E)**. In the spleen, WNV RNA was detected as early as 2 dpi and remained elevated through 9 dpi, by which time viral RNA levels were higher in the CNS **(Fig. 1F)**. In contrast, WNV RNA was largely undetectable in the liver, with sporadic detection at low levels in individual animals **(Fig. 1G)**. Thus, WNV infection in mice on a mixed C57BL/6J × 129S genetic background followed the expected progression from peripheral infection to widespread CNS dissemination, consistent with the kinetics previously reported in C57BL/6J mice [50].

**Figure 1.**
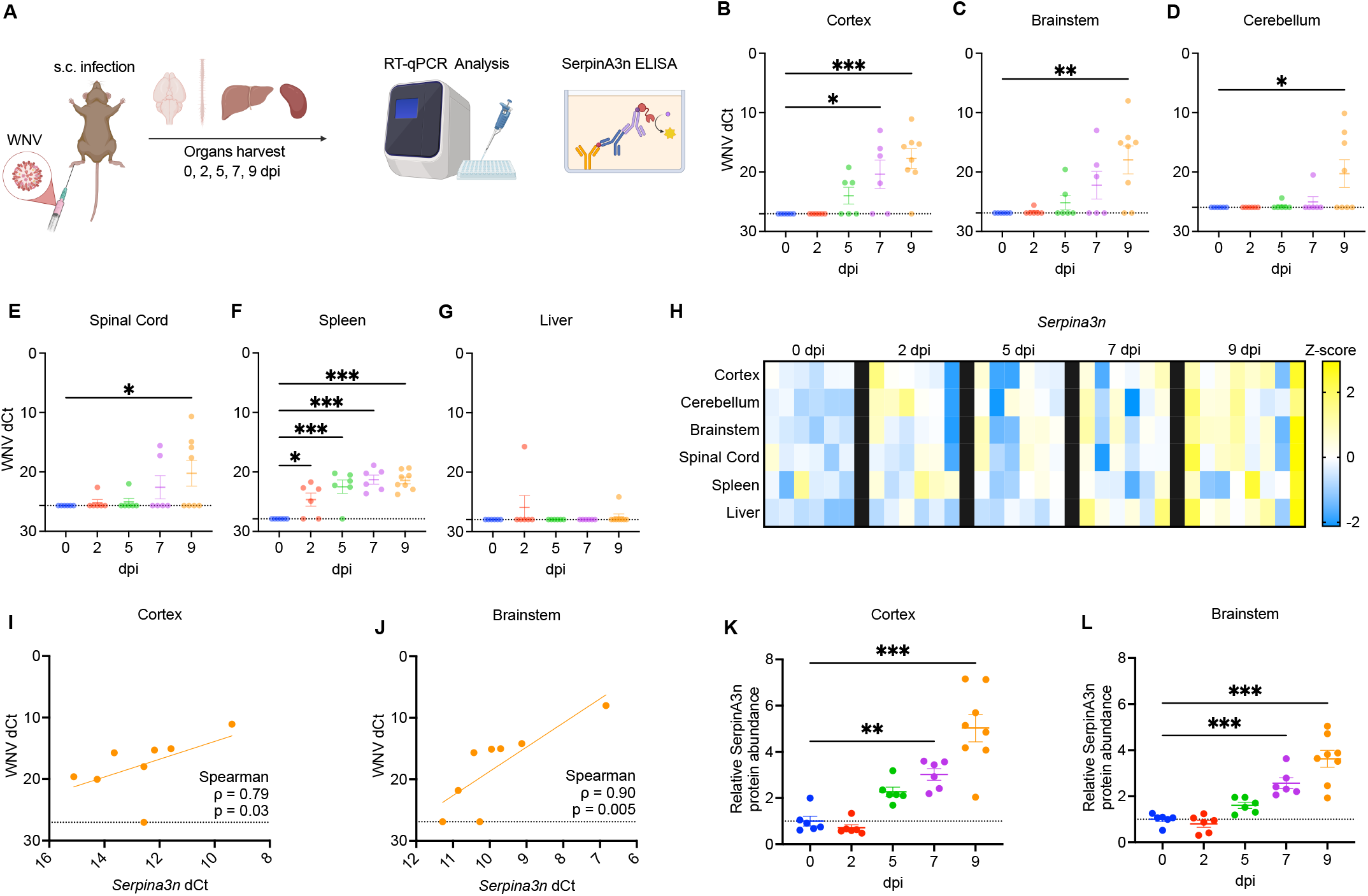
Subcutaneous WNV infection in mixed C57BL/6J × 129S mice results in CNS dissemination and induction of SerpinA3n expression. (A) Experimental schematic, created with BioRender. Mice were inoculated subcutaneously in the footpad with 1,875 PFU of WNV WN02-Bird114. Cortex, brainstem, cerebellum, spinal cord, spleen, and liver were harvested after perfusion on indicated days post-infection (dpi) for RT-qPCR analysis of WNV RNA and *Serpina3n* transcripts and for SerpinA3n ELISA. (B–G) WNV RNA levels (WNV dCt, normalized to *18S*) in the cortex (B), brainstem (C), cerebellum (D), spinal cord (E), spleen (F), and liver (G) at the indicated dpi. The dotted line marks the limit of detection. (H) Heatmap depicting *Serpina3n* transcript levels across tissues and time points. Z-scores were calculated from ddCt values. Each square represents an individual mouse. (I, J) Correlation between *Serpina3n* dCt and WNV dCt in individual mice at 9 dpi in the cortex (I) and brainstem (J). Spearman correlation coefficients (ρ) and p-values are shown. The linear regression line is shown for visualization only. The dotted line marks the limit of detection for WNV RNA. (K, L) Relative SerpinA3n protein abundance in the cortex (K) and brainstem (L), measured by ELISA and normalized to the 0 dpi group (dotted line). Each dot represents an individual mouse. Data are presented as mean ± SEM (B–G, K, L). For B–G, K, and L, each group was compared with the 0 dpi control group by one-way ANOVA with Dunnett’s multiple comparisons test. Only significant comparisons are shown. *p < 0.05, **p < 0.01, ***p < 0.001. n = 6 mice per group at 0, 2, 5, and 7 dpi and n = 8 mice at 9 dpi.

Given that SerpinA3n is an acute-phase protein induced by inflammatory stimuli [30], we next examined the kinetics and tissue distribution of *Serpina3n* expression during WNV infection. *Serpina3n* transcript levels were significantly elevated at 9 dpi in the cortex, cerebellum, brainstem, and spinal cord **(Fig. 1H)**, coinciding with increased WNV RNA levels in these regions **(Fig. 1B–E)**. *Serpina3n* expression was also elevated in the spleen and liver at 9 dpi **(Fig. 1H)**, with induction in the liver evident as early as 7 dpi despite the near absence of detectable viral RNA **(Fig. 1G)**. Several other serpin family members (*Serpina3g*, *Serpina3h*, *Serpinb8*, *Serpine1*, and *Serping1*) were similarly upregulated at 9 dpi in the cortex, brainstem, spinal cord, and liver, with more variable changes in the cerebellum and spleen **(Fig. S1A–F)**.

We next assessed the relationship between *Serpina3n* expression and viral burden in individual CNS regions at 9 dpi. *Serpina3n* transcript levels correlated positively with WNV RNA levels in the cortex (Spearman ρ = 0.79, *p* = 0.03), brainstem (ρ = 0.90, *p* = 0.005), and spinal cord (ρ = 0.82, *p* = 0.02) **(Fig. 1I, J; Fig. S1G)**, but not in the cerebellum (ρ = 0.46, *p* = 0.26; **Fig. S1H**). Induction of *Serpina3n* expression was also evident at the protein level, with significantly elevated SerpinA3n protein concentrations in both the cortex and brainstem at 7 and 9 dpi **(Fig. 1K, L)**. Together, these findings demonstrate that neuroinvasive WNV infection is accompanied by widespread induction of serpin expression, including substantial increases in SerpinA3n protein abundance in the infected brain.

### SerpinA3n activity in the CNS promotes survival and preserves BBB integrity during WNV encephalitis

Given the induction of *Serpina3n* expression in both peripheral tissues and the CNS of WNV-infected mice **(Fig. 1)**, we sought to determine the relative contributions of SerpinA3n in peripheral and CNS compartments to host protection during infection. We first administered the SerpinA3n inhibitor ARN1468 (2.7 mg/kg, i.p.) twice daily from 4 to 8 dpi, a window encompassing neuroinvasion and the onset of clinical disease but preceding substantial mortality **(Fig. 2A)**. Peripheral administration of ARN1468 had no significant effect on survival relative to vehicle treatment **(Fig. 2B)**. Given the poor brain penetrance of ARN1468 following peripheral administration [51], we next administered ARN1468 directly into the CNS (5 µg, i.c.v.) at 5 dpi **(Fig. 2C)**. Localized inhibition of SerpinA3n within the CNS accelerated the onset of mortality and significantly reduced survival, with only 9% of ARN1468-treated mice surviving to 21 dpi compared with 60% of vehicle-treated mice **(Fig. 2D)**. All mock-infected mice receiving i.c.v. ARN1468 survived **(Fig. 2D)**, suggesting that this effect was not due to generalized toxicity associated with ARN1468 delivery to the CNS.

**Figure 2.**
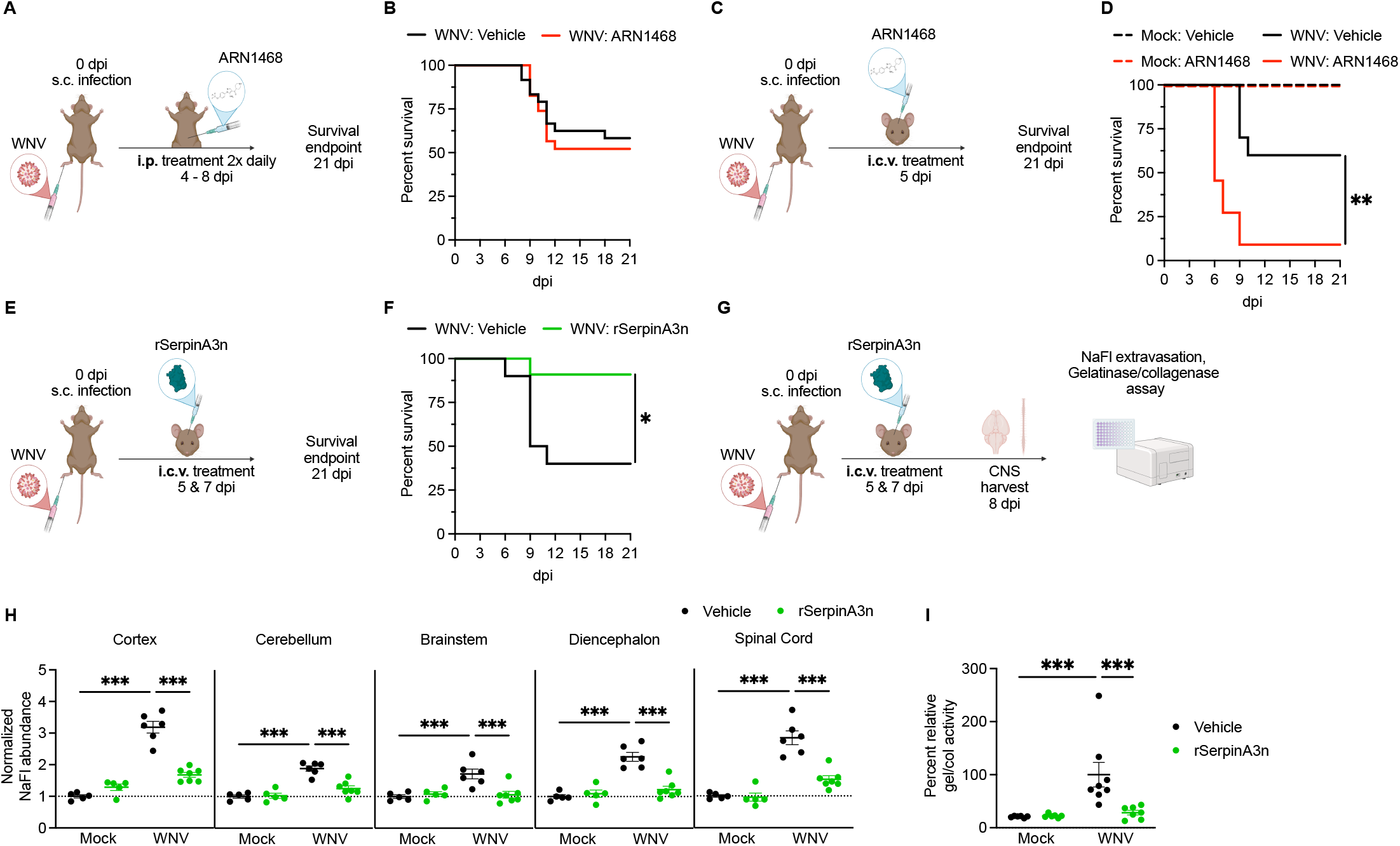
SerpinA3n activity in the CNS promotes survival and preserves BBB integrity during WNV encephalitis. (A) Schematic of systemic SerpinA3n inhibition, created with BioRender. WNV-infected mice were treated intraperitoneally (i.p.) with the SerpinA3n inhibitor ARN1468 (2.7 mg/kg) twice daily from 4 to 8 days post-infection (dpi) and monitored for survival through 21 dpi. (B) Survival of WNV-infected mice treated i.p. with vehicle or ARN1468. (C) Schematic of CNS-localized SerpinA3n inhibition. WNV-infected mice received a single intracerebroventricular (i.c.v.) injection of ARN1468 (5 µg) at 5 dpi and were monitored for survival through 21 dpi. (D) Survival of mock- and WNV-infected mice treated i.c.v. with vehicle or ARN1468. (E) Schematic of recombinant SerpinA3n (rSerpinA3n) supplementation. WNV-infected mice received i.c.v. rSerpinA3n (4 ng) or vehicle at 5 and 7 dpi and were monitored for survival through 21 dpi. (F) Survival of WNV-infected mice treated i.c.v. with vehicle or rSerpinA3n. (G) Schematic for assessing BBB integrity and proteolytic activity. Mock- and WNV-infected mice received i.c.v. vehicle or rSerpinA3n at 5 and 7 dpi, and CNS tissues were harvested at 8 dpi. (H) BBB permeability, measured as sodium fluorescein (NaFl) abundance in indicated CNS regions, normalized to the Mock:Vehicle group. The dotted line marks the Mock:Vehicle baseline (value of 1). (I) Percent relative gelatinase/collagenase (gel/col) activity in the CNS, normalized relative to the WNV:Vehicle group. In H and I, each dot represents an individual mouse. Data are presented as mean ± SEM. Survival curves in B, D, and F were compared by log-rank (Mantel-Cox) test; in D, all four groups were compared. For H and I, a two-way ANOVA was performed, followed by uncorrected Fisher’s LSD test for three comparisons (Mock:Vehicle vs. WNV:Vehicle, Mock:Vehicle vs. Mock:rSerpinA3n, and WNV:Vehicle vs. WNV:rSerpinA3n), made within each CNS region in H. Only significant comparisons are shown. *p < 0.05, **p < 0.01, ***p < 0.001. Sample sizes: (B) WNV:Vehicle n = 24, WNV:ARN1468 n = 23. (D) WNV:Vehicle n = 10, WNV:ARN1468 n = 11, Mock:Vehicle n = 10, Mock:ARN1468 n = 10. (F) WNV:Vehicle n = 10, WNV:rSerpinA3n n = 11. (H) Mock:Vehicle n = 5, Mock:rSerpinA3n n = 5, WNV:Vehicle n = 6, WNV:rSerpinA3n n = 7. (I) Mock:Vehicle n = 6, Mock:rSerpinA3n n = 6, WNV:Vehicle n = 8, WNV:rSerpinA3n n = 7.

As a complementary approach, we also administered recombinant SerpinA3n (rSerpinA3n; 4 ng, i.c.v.) to WNV-infected mice at 5 and 7 dpi **(Fig. 2E)**. CNS supplementation with rSerpinA3n significantly improved survival, with 91% of treated mice surviving to 21 dpi compared with 40% of vehicle-treated mice **(Fig. 2F)**. To investigate the basis for this protection, we assessed BBB permeability and proteolytic activity in the CNS at 8 dpi **(Fig. 2G)**. BBB permeability was assessed by quantifying the extravasation of sodium fluorescein, a small-molecule fluorescent tracer that is largely excluded from the CNS by an intact BBB [50,52]. WNV infection significantly increased the accumulation of sodium fluorescein in all CNS regions examined, indicative of widespread BBB disruption. Treatment with rSerpinA3n significantly reduced sodium fluorescein accumulation in the cortex, cerebellum, brainstem, diencephalon, and spinal cord, restoring levels toward those observed in mock-infected mice **(Fig. 2H)**.

Gelatinases and collagenases, including matrix metalloproteinases (MMPs) such as MMP-2 and MMP-9, contribute to BBB disruption in the inflamed CNS through the degradation of basement membrane and junctional proteins [53]. We therefore asked whether the preservation of BBB integrity by rSerpinA3n was accompanied by reduced activity of these proteases. WNV infection significantly increased gelatinase/collagenase activity in the brain, whereas treatment with rSerpinA3n reduced this activity to near-baseline levels **(Fig. 2I)**. CNS viral burden at 8 dpi did not differ significantly between rSerpinA3n- and vehicle-treated mice **(Fig. S2)**. Together, these findings demonstrate that SerpinA3n activity within the CNS promotes host survival during WNV encephalitis and that this protection is associated with the preservation of BBB integrity and suppression of proteolytic activity rather than changes in CNS viral burden.

### CNS delivery of rSerpinA3n limits leukocyte recruitment and antigen-presenting cell activation during WNV encephalitis

Given that rSerpinA3n treatment improved survival and preserved BBB integrity during WNV encephalitis, we next asked whether SerpinA3n influences leukocyte accumulation and activation in the infected CNS. Following i.c.v. administration of rSerpinA3n, brains were harvested at 8 dpi, and CNS leukocyte populations were characterized by flow cytometry **(Fig. 3A)**. Treatment with rSerpinA3n significantly reduced the numbers of both infiltrating (CD45.2^hi^) and CNS-resident (CD45.2^int^) immune cells relative to vehicle treatment **(Fig. 3B)**. Numbers of both CD3⁺ T cells and CD19⁺ B cells were significantly reduced **(Fig. 3C)**. Within the T cell compartment, rSerpinA3n treatment reduced the numbers of both CD4⁺ and CD8α⁺ T cells **(Fig. 3D, E)** without altering their relative proportions among CD3⁺ T cells **(Fig. 3D, F)**, indicating a reduction in overall T cell accumulation rather than a preferential reduction in either subset.

**Figure 3.**
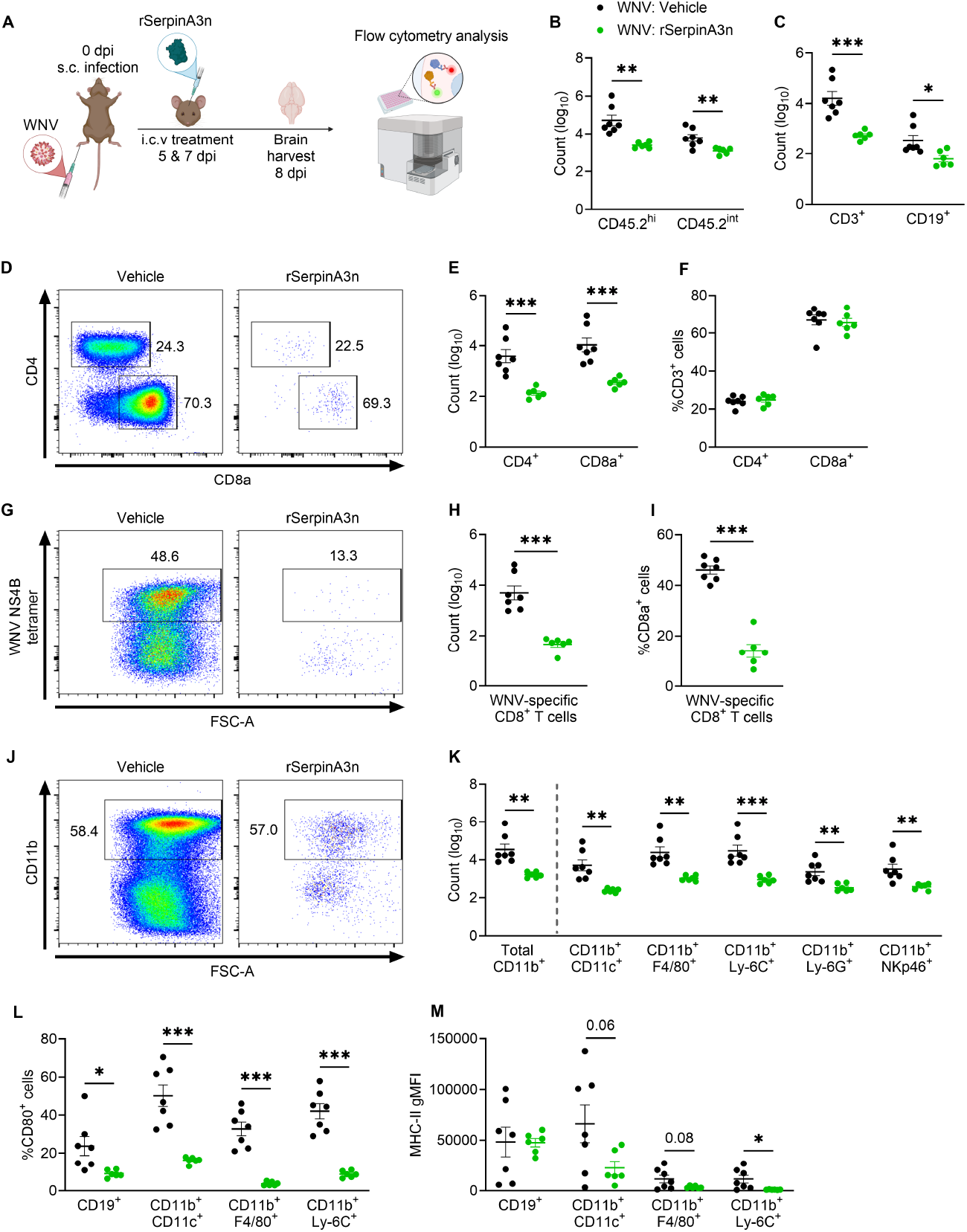
CNS delivery of rSerpinA3n limits leukocyte recruitment and antigen-presenting cell activation during WNV encephalitis. (A) Schematic of the experiment, created with BioRender. WNV-infected mice received intracerebroventricular (i.c.v.) vehicle or recombinant SerpinA3n (rSerpinA3n) at 5 and 7 days post-infection (dpi). Brains were harvested at 8 dpi, and CNS leukocytes were analyzed by flow cytometry. (B) Numbers of infiltrating (CD45.2^hi^) and resident (CD45.2^int^) immune cells, gated on single live cells. (C) Numbers of CD3^+^ T cells and CD19^+^ B cells, gated on CD45.2^hi^ cells. (D) Representative flow cytometry plots of CD4 versus CD8α expression, gated on CD3^+^ cells; numbers indicate the percentage of cells in each gate. (E, F) Numbers (E) and percentages among CD3^+^ cells (F) of CD4^+^ and CD8α^+^ T cells. (G) Representative flow cytometry plots of WNV NS4B tetramer staining versus FSC-A, gated on CD8α^+^ cells; numbers indicate the percentage of tetramer^+^ cells. (H, I) Number of WNV-specific (NS4B tetramer^+^) CD8α^+^ T cells (H) and their percentage among CD8α^+^ cells (I). (J) Representative flow cytometry plots of CD11b versus FSC-A, gated on CD45.2^hi^ cells; numbers indicate the percentage of CD11b^+^ cells. (K) Numbers of total CD11b^+^ cells (gated on CD45.2^hi^ cells) and of the indicated populations: CD11b^+^CD11c^+^, CD11b^+^F4/80^+^, CD11b^+^Ly-6C^+^, CD11b^+^Ly-6G^+^, and CD11b^+^NKp46^+^ cells (each gated on CD11b^+^ cells). (L) Percentage of CD80^+^ cells among CD19^+^ B cells (gated on CD45.2^hi^ cells) and among CD11b^+^CD11c^+^, CD11b^+^F4/80^+^, and CD11b^+^Ly-6C^+^ cells (each gated on CD11b^+^ cells). (M) MHC-II geometric mean fluorescence intensity (gMFI) in the same populations as in L. In B, C, E, F, H, I, and K, cell counts are shown on a log_10_ scale. Each dot represents an individual mouse. Data are presented as mean ± SEM. Vehicle and rSerpinA3n groups were compared within each cell population by Student’s unpaired t tests. Exact p-values are shown for non-significant comparisons. *p < 0.05, **p < 0.01, ***p < 0.001. n = 7 for WNV:Vehicle and n = 6 for WNV:rSerpinA3n.

To assess the accumulation of virus-specific T cells in the CNS, we identified WNV-specific CD8⁺ T cells via labeling with an H-2Dᵇ/NS4B tetramer **(Fig. 3G–I)**. Treatment with rSerpinA3n markedly reduced both the number of WNV-specific CD8⁺ T cells **(Fig. 3G, H)** and their frequency among CD8⁺ T cells **(Fig. 3G, I)**. Given that the proportion of CD8⁺ cells among CD3⁺ T cells was unchanged **(Fig. 3F)**, these data suggest that rSerpinA3n treatment disproportionately reduced the accumulation of virus-specific CD8⁺ T cells. This is notable given the established contributions of WNV-specific CD8⁺ T cells, including those recognizing the immunodominant NS4B epitope, to host survival [54,55] and viral clearance from the CNS [42]. The reduced frequency of WNV-specific CD8⁺ T cells may reflect changes in their recruitment or expansion within the CNS.

Among innate immune cell populations, total CD11b⁺ cell numbers were significantly reduced following rSerpinA3n treatment **(Fig. 3J, K)**. This reduction extended to CD11b⁺CD11c⁺ dendritic cells, CD11b⁺F4/80⁺ macrophages, CD11b⁺Ly-6C⁺ monocytes, CD11b⁺Ly-6G⁺ neutrophils, and CD11b⁺NKp46⁺ natural killer cells **(Fig. 3K)**. To assess the activation state of antigen-presenting cells (APCs), we examined expression of CD80 and MHC-II. The frequency of CD80⁺ cells was significantly reduced across all APC populations examined, including B cells, dendritic cells, macrophages, and monocytes **(Fig. 3L)**. MHC-II expression, measured by geometric mean fluorescence intensity (gMFI), was significantly reduced on CD11b⁺Ly-6C⁺ monocytes, with trends toward reduced expression on CD11b⁺CD11c⁺ dendritic cells (*p* = 0.06) and CD11b⁺F4/80⁺ macrophages (*p* = 0.08) **(Fig. 3M)**.

To determine whether these effects were dependent on infection, we performed the same analyses in mock-infected mice. In the absence of infection, rSerpinA3n treatment had minimal effects on CNS leukocyte numbers and did not significantly alter CD80 or MHC-II expression on APCs **(Fig. S3A–F)**. The only significant differences were a modest increase in CD8α⁺ T cell numbers and a modest decrease in the proportion of CD4⁺ cells among CD3⁺ T cells **(Fig. S3B, C)**, indicating that the broader immunomodulatory effects of rSerpinA3n were dependent on infection.

Together, these data demonstrate that the protective effects of rSerpinA3n during WNV encephalitis are accompanied by reduced CNS leukocyte accumulation and APC activation. Notably, despite a reduction in the abundance of virus-specific CD8⁺ T cells, rSerpinA3n treatment did not measurably affect CNS viral burden **(Fig. S2)**. These findings suggest that attenuation of CNS inflammation may contribute to the protective effects of SerpinA3n treatment, although whether SerpinA3n directly regulates leukocyte recruitment or does so indirectly through the preservation of BBB integrity remains to be established.

### Loss of either astrocytic or neuronal SerpinA3n exacerbates WNV-induced mortality, BBB disruption, and proteolytic activity in the CNS

Having established a protective role for CNS-localized SerpinA3n through pharmacological inhibition and recombinant protein supplementation, we next investigated the contributions of endogenous SerpinA3n production by resident CNS cells to the regulation of WNV pathogenesis. Because SerpinA3n is expressed by both astrocytes and neurons [30], we generated mice selectively deficient in astrocytic or neuronal SerpinA3n by crossing mice harboring loxP sites flanking exon 2 of the endogenous *Serpina3n* locus [56] (*Serpina3n*^fl/fl^) to mice expressing either tamoxifen-inducible Cre recombinase under the control of the *Aldh1l1* promoter or constitutive Cre recombinase under the control of the *Syn1* promoter. To assess the contributions of astrocytes and neurons to SerpinA3n production during infection, we quantified SerpinA3n protein levels in brains harvested at 0, 5, and 8 dpi by ELISA **(Fig. 4A)**. Although brain SerpinA3n levels did not differ at 0 dpi, mice lacking astrocytic or neuronal SerpinA3n (*Serpina3n*^fl/fl^ *Aldh1l1*-CreERT2⁺ and *Serpina3n*^fl/fl^ *Syn1*-Cre⁺ mice, respectively) exhibited significantly reduced SerpinA3n levels at 5 and 8 dpi relative to their respective Cre-negative littermate controls **(Fig. 4B, C)**. By 8 dpi, SerpinA3n protein levels were reduced by approximately half or more in both lines, identifying astrocytes and neurons as major sources of SerpinA3n in the WNV-infected CNS.

**Figure 4.**
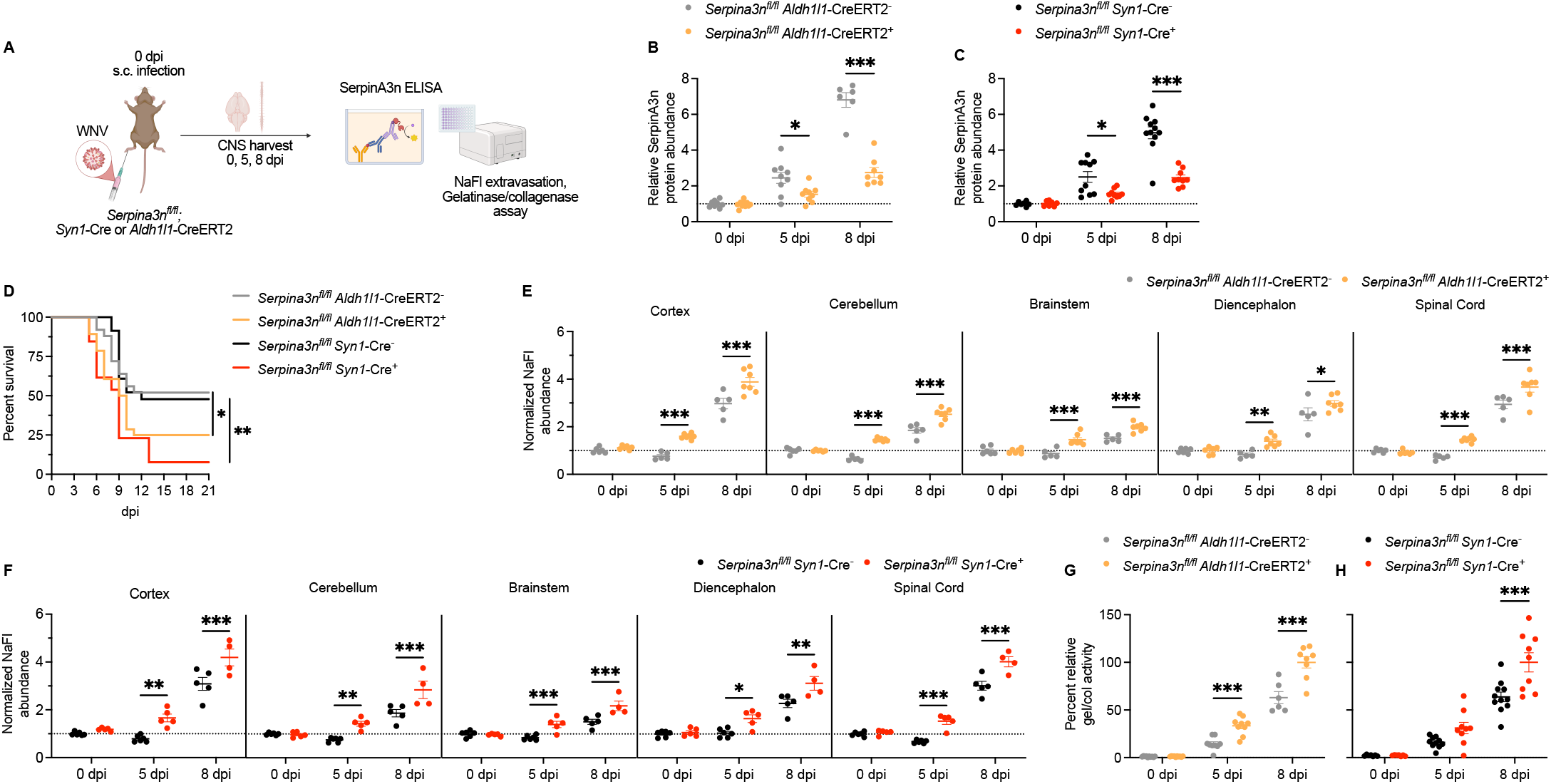
Loss of either astrocytic or neuronal SerpinA3n exacerbates WNV-induced mortality, BBB disruption, and proteolytic activity in the CNS. (A) Schematic of the experiment, created with BioRender. *Serpina3n^fl/fl^ Aldh1l1*-CreERT2^+^ (astrocytic deletion) and *Serpina3n^fl/fl^ Syn1*-Cre^+^ (neuronal deletion) mice and their Cre-negative littermate controls were infected subcutaneously with WNV. CNS tissues were harvested at 0, 5, and 8 days post-infection (dpi) for indicated assays. (B, C) Relative SerpinA3n protein abundance in the CNS of *Serpina3n^fl/fl^ Aldh1l1*-CreERT2 (B) and *Serpina3n^fl/fl^ Syn1*-Cre (C) mice, measured by ELISA. (D) Survival of *Serpina3n^fl/fl^ Aldh1l1*-CreERT2^+^ and *Serpina3n^fl/fl^ Syn1*-Cre^+^ mice and their Cre-negative controls. (E, F) BBB permeability, measured as normalized sodium fluorescein (NaFl) abundance in indicated CNS regions of *Serpina3n^fl/fl^ Aldh1l1*-CreERT2 (E) and *Serpina3n^fl/fl^ Syn1*-Cre (F) mice. (G, H) Percent relative gelatinase/collagenase (gel/col) activity in the CNS of *Serpina3n^fl/fl^ Aldh1l1*-CreERT2 (G) and *Serpina3n^fl/fl^ Syn1*-Cre (H) mice, expressed as a percentage of the activity in the 8 dpi Cre-positive group of the same genotype. In B, C, E, and F, the dotted line marks normalization to the 0 dpi Cre-negative control group of the respective genotype. In B, C, and E–H, each dot represents an individual mouse. Data are presented as mean ± SEM. For B, C, E, F, G, and H, a two-way ANOVA was performed with Šídák’s multiple comparisons test, comparing Cre-negative and Cre-positive groups within each dpi. In D, survival was compared by log-rank (Mantel-Cox) test, with Cre-positive mice compared only with Cre-negative mice of the same genotype. Only significant comparisons are shown. *p < 0.05, **p < 0.01, ***p < 0.001. Sample sizes (Cre-negative, Cre-positive): (B, G) *Serpina3n^fl/fl^ Aldh1l1*-CreERT2: 0 dpi: 10 and 10; 5 dpi: 9 and 9; 8 dpi: 6 and 8. (C, H) *Serpina3n^fl/fl^ Syn1*-Cre: 0 dpi: 7 and 7; 5 dpi: 10 and 9; 8 dpi: 11 and 9. (D) *Serpina3n^fl/fl^ Aldh1l1*-CreERT2: 25 and 28; *Serpina3n^fl/fl^ Syn1*-Cre: 23 and 13. (E) *Serpina3n^fl/fl^ Aldh1l1*-CreERT2: 0 dpi: 6 and 6; 5 dpi: 5 and 7; 8 dpi: 5 and 7. (F) *Serpina3n^fl/fl^ Syn1*-Cre: 0 dpi: 6 and 5; 5 dpi: 6 and 5; 8 dpi: 5 and 4.

To determine whether SerpinA3n production by either cell population was protective during WNV encephalitis, we challenged these mice with WNV and monitored survival. Both *Serpina3n*^fl/fl^ *Aldh1l1*-CreERT2⁺ and *Serpina3n*^fl/fl^ *Syn1*-Cre⁺ mice exhibited significantly increased mortality relative to their Cre-negative littermate controls **(Fig. 4D)**. In contrast, we observed no evidence of increased susceptibility to WNV infection in heterozygous *Serpina3n*^fl/+^ *Aldh1l1*-CreERT2⁺ or *Serpina3n*^fl/+^ *Syn1*-Cre⁺ mice **(Fig. S4A)**. Assessment of BBB permeability revealed significantly increased sodium fluorescein extravasation in both *Serpina3n*^fl/fl^ *Aldh1l1*-CreERT2⁺ and *Serpina3n*^fl/fl^ *Syn1*-Cre⁺ mice at 5 and 8 dpi across all CNS regions examined (cortex, cerebellum, brainstem, diencephalon, and spinal cord), with no differences observed at 0 dpi **(Fig. 4E, F)**.

Mice lacking astrocytic SerpinA3n also exhibited significantly increased gelatinase/collagenase activity in the CNS at 5 and 8 dpi relative to their Cre-negative littermate controls **(Fig. 4G)**. Similarly, mice lacking neuronal SerpinA3n exhibited significantly increased gelatinase/collagenase activity at 8 dpi **(Fig. 4H)**. These findings complement our observation that CNS supplementation of rSerpinA3n reduced gelatinase/collagenase activity during WNV infection **(Fig. 2I)**. Neither astrocytic nor neuronal SerpinA3n deficiency significantly affected CNS viral burden at 5 or 8 dpi **(Fig. S4B, C)**, suggesting that the increased susceptibility of these mice to WNV-induced mortality was not attributable to increased viral burden. Thus, loss of either astrocytic or neuronal SerpinA3n exacerbated mortality, BBB disruption, and proteolytic activity during WNV encephalitis, mirroring the protective effects of CNS supplementation with rSerpinA3n on BBB integrity and proteolytic activity **(Fig. 2H, I)**. Collectively, these data identify both astrocytes and neurons as important contributors to the endogenous regulation of proteolysis and maintenance of neurovascular integrity during CNS viral infection.

### Loss of neuronal, but not astrocytic, SerpinA3n exacerbates CNS leukocyte recruitment during WNV encephalitis

Given that loss of either astrocytic or neuronal SerpinA3n exacerbated mortality, BBB disruption, and proteolytic activity during WNV encephalitis, we next asked whether either cellular source of SerpinA3n regulates CNS leukocyte recruitment. We characterized CNS immune cell populations in mice lacking astrocytic or neuronal SerpinA3n by flow cytometry **(Fig. 5A)**. Numbers of both infiltrating (CD45.2^hi^) and CNS-resident (CD45.2^int^) immune cells were significantly increased in *Serpina3n*^fl/fl^ *Syn1*-Cre⁺ mice relative to their Cre-negative littermate controls, whereas no significant differences were observed in *Serpina3n*^fl/fl^ *Aldh1l1*-CreERT2⁺ mice **(Fig. 5B)**. Among adaptive immune cell populations, numbers of CD3⁺ T cells, including both CD4⁺ and CD8α⁺ T cells, were significantly increased in mice lacking neuronal, but not astrocytic, SerpinA3n **(Fig. 5C, D)**. Notably, the proportion of CD8α⁺ cells among CD3⁺ T cells was modestly increased in mice lacking astrocytic SerpinA3n but was unchanged in mice lacking neuronal SerpinA3n **(Fig. 5E)**. Loss of neuronal, but not astrocytic, SerpinA3n resulted in significantly increased numbers of WNV-specific CD8α⁺ T cells **(Fig. 5F)**, with a trend toward increased frequency of WNV-specific cells among CD8α⁺ T cells (*p* = 0.08) **(Fig. 5G)**.

**Figure 5.**
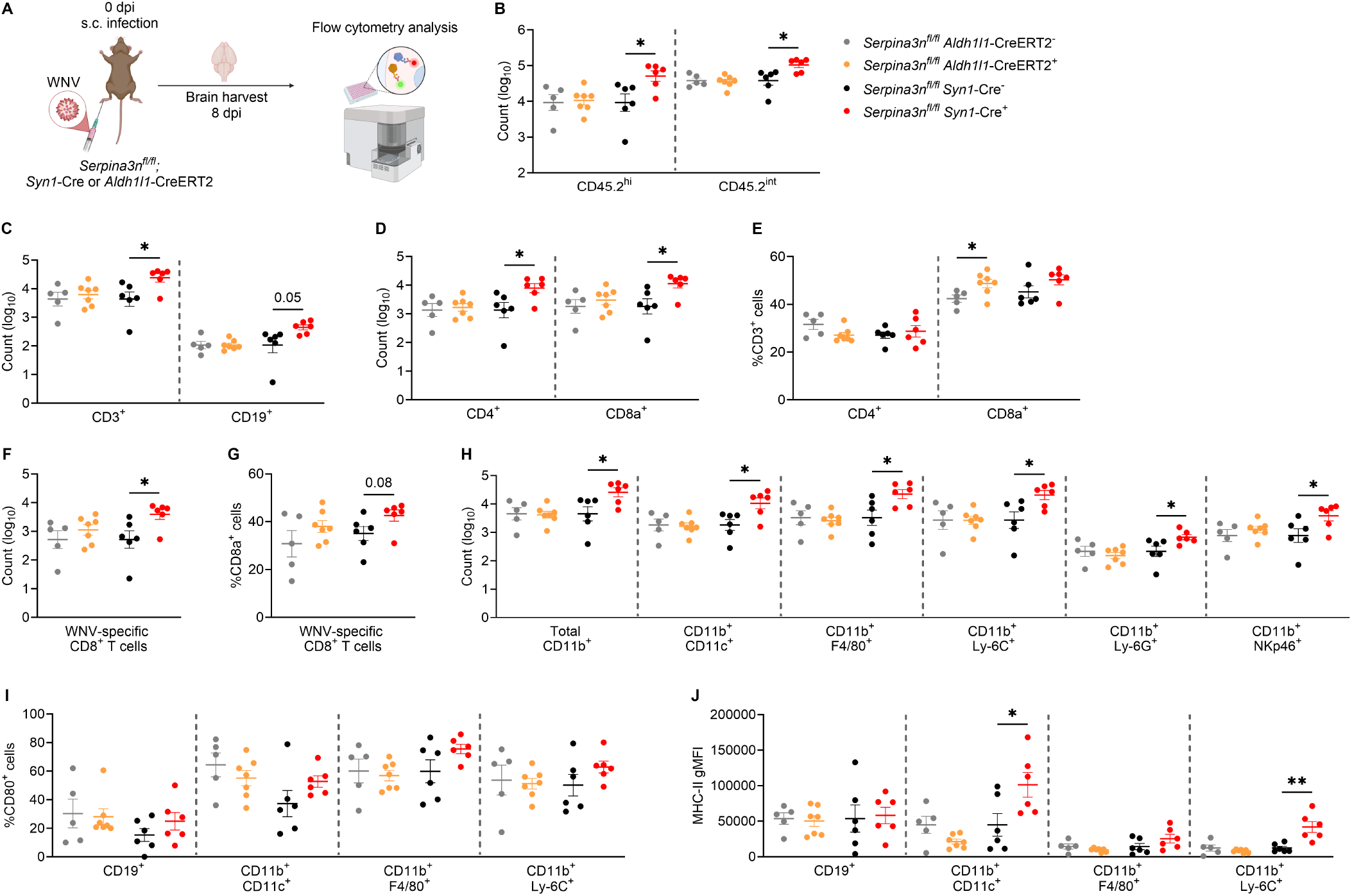
Loss of neuronal, but not astrocytic, SerpinA3n exacerbates CNS leukocyte recruitment during WNV encephalitis. (A) Schematic of the experiment, created with BioRender. *Serpina3n^fl/fl^ Aldh1l1*-CreERT2^+^ and *Serpina3n^fl/fl^ Syn1*-Cre^+^ mice and their Cre-negative littermate controls were infected subcutaneously with WNV. Brains were harvested at 8 days post-infection (dpi), and CNS leukocytes were analyzed by flow cytometry. (B) Numbers of infiltrating (CD45.2^hi^) and resident (CD45.2^int^) immune cells, gated on live single cells. (C) Numbers of CD3^+^ T cells and CD19^+^ B cells, gated on CD45.2^hi^ cells. (D, E) Numbers (D) and percentages among CD3^+^ cells (E) of CD4^+^ and CD8α^+^ T cells, gated on CD3^+^ cells. (F, G) Number of WNV-specific (NS4B tetramer^+^) CD8α^+^ T cells (F) and their percentage among CD8α^+^ cells (G). (H) Numbers of total CD11b^+^ cells (gated on CD45.2^hi^ cells) and of the indicated populations: CD11b^+^CD11c^+^, CD11b^+^F4/80^+^, CD11b^+^Ly-6C^+^, CD11b^+^Ly-6G^+^, and CD11b^+^NKp46^+^ cells (each gated on CD11b^+^ cells). (I) Percentage of CD80^+^ cells among CD19^+^ B cells (gated on CD45.2^hi^ cells) and among CD11b^+^CD11c^+^, CD11b^+^F4/80^+^, and CD11b^+^Ly-6C^+^ cells (each gated on CD11b^+^ cells). (J) MHC-II geometric mean fluorescence intensity (gMFI) in the same populations as in I. In B, C, D, F, and H, cell counts are shown on a log_10_ scale. Each dot represents an individual mouse. Data are presented as mean ± SEM. Within each genotype, Cre-negative and Cre-positive groups were compared within each cell population by Student’s unpaired t tests. Exact p-values are shown for non-significant comparisons in C and G. *p < 0.05, **p < 0.01. n = 5 for *Serpina3n^fl/fl^ Aldh1l1*-CreERT2^-^, n = 7 for *Serpina3n^fl/fl^ Aldh1l1*-CreERT2^+^, n = 6 for *Serpina3n^fl/fl^ Syn1*-Cre^-^, and n = 6 for *Serpina3n^fl/fl^ Syn1*-Cre^+^.

Similarly, numbers of total CD11b⁺ cells, CD11b⁺CD11c⁺ dendritic cells, CD11b⁺F4/80⁺ macrophages, CD11b⁺Ly-6C⁺ monocytes, CD11b⁺Ly-6G⁺ neutrophils, and CD11b⁺NKp46⁺ natural killer cells were significantly increased in mice lacking neuronal, but not astrocytic, SerpinA3n **(Fig. 5H)**. The frequency of CD80⁺ cells among individual APC populations was unaffected by loss of either neuronal or astrocytic SerpinA3n **(Fig. 5I)**. However, MHC-II expression was significantly elevated on CD11b⁺CD11c⁺ dendritic cells and CD11b⁺Ly-6C⁺ monocytes in mice lacking neuronal SerpinA3n **(Fig. 5J)**.

Thus, the increased susceptibility of mice lacking neuronal SerpinA3n to WNV-induced mortality was accompanied by increased accumulation of innate and adaptive immune cells in the CNS and elevated MHC-II expression on select APC populations, complementing our observations in mice treated with rSerpinA3n **(Fig. 3)**. In contrast, loss of astrocytic SerpinA3n had little effect on CNS leukocyte abundance or APC activation despite increased BBB permeability and gelatinase/collagenase activity **(Fig. 4E, G)**. These findings suggest that, although both astrocytic and neuronal SerpinA3n contribute to the regulation of CNS proteolysis and preservation of BBB integrity, the immunoregulatory functions of SerpinA3n may differ depending on the cellular source and spatial context of its production.

### SerpinA3n restrains pathogenic granzyme B activity in the infected CNS

Given that SerpinA3n is a potent inhibitor of granzyme B [27,29], we asked whether regulation of granzyme B activity contributes to the protective effects of SerpinA3n during WNV encephalitis. WNV infection significantly increased granzyme B activity in the CNS relative to mock-infected mice, and CNS administration of rSerpinA3n significantly, albeit partially, attenuated this increase **(Fig. 6A)**. Conversely, loss of either astrocytic or neuronal SerpinA3n resulted in significantly increased granzyme B activity in the CNS at both 5 and 8 dpi relative to Cre-negative littermate controls, with no differences observed at 0 dpi **(Fig. 6B, C)**. These findings demonstrate that SerpinA3n produced by both astrocytes and neurons contributes to the endogenous regulation of granzyme B activity during WNV encephalitis.

**Figure 6.**
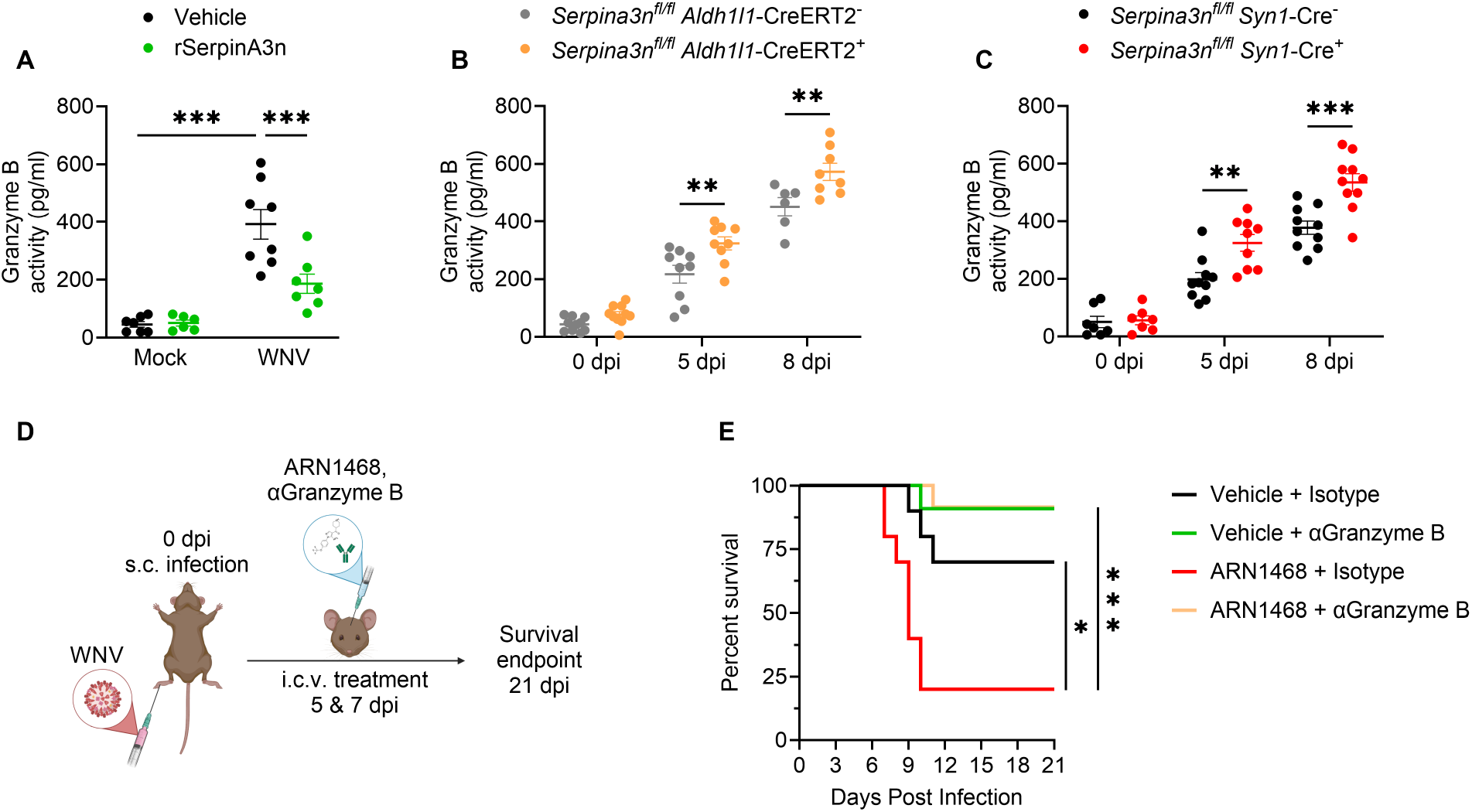
SerpinA3n restrains pathogenic granzyme B activity in the infected CNS. (A) Granzyme B activity, measured with an immunocapture enzymatic activity assay, in the CNS of mock- and WNV-infected mice at 8 days post-infection (dpi) treated intracerebroventricularly (i.c.v.) with vehicle or recombinant SerpinA3n (rSerpinA3n). (B, C) Granzyme B activity in the CNS of *Serpina3n^fl/fl^ Aldh1l1*-CreERT2^+^ (B) and *Serpina3n^fl/fl^ Syn1*-Cre^+^ (C) mice and their Cre-negative littermate controls at 0, 5, and 8 dpi. (D) Schematic of the Granzyme B neutralization experiment, created with BioRender. WNV-infected mice received i.c.v. treatment with the SerpinA3n inhibitor ARN1468 or vehicle, together with an anti-Granzyme B neutralizing antibody (αGranzyme B) or isotype control, at 5 and 7 dpi and were monitored for survival through 21 dpi. (E) Survival of WNV-infected mice receiving indicated treatments. In A–C, each dot represents an individual mouse, and data are presented as mean ± SEM. For A-C, comparisons were made using two-way ANOVA with Šídák’s multiple comparisons test. In E, survival was compared by log-rank (Mantel-Cox) test. Only significant comparisons are shown. *p < 0.05, **p < 0.01, ***p < 0.001. Sample sizes: (A) Mock:Vehicle: n = 6; Mock:rSerpinA3n: n = 6; WNV:Vehicle: n = 8; WNV:rSerpinA3n: n = 7. (B) *Serpina3n^fl/fl^ Aldh1l1*-CreERT2 (Cre-negative, Cre-positive): 0 dpi: 10 and 10; 5 dpi: 9 and 9; 8 dpi: 6 and 8. (C) *Serpina3n^fl/fl^ Syn1*-Cre (Cre-negative, Cre-positive): 0 dpi: 7 and 7; 5 dpi: 10 and 9; 8 dpi: 11 and 9. (E) Vehicle + Isotype: n = 10; Vehicle + αGranzyme B: n = 11; ARN1468 + Isotype: n = 10; ARN1468 + αGranzyme B: n = 12.

To determine whether granzyme B activity contributes to the increased mortality associated with CNS pharmacological inhibition of SerpinA3n, we i.c.v. treated WNV-infected wild-type mice with ARN1468 in combination with a granzyme B-neutralizing antibody or an isotype control antibody **(Fig. 6D)**. Consistent with our earlier findings, SerpinA3n inhibition significantly reduced survival in mice receiving the isotype control antibody, with 20% of ARN1468-treated mice surviving compared with 70% of vehicle-treated mice. Notably, neutralization of granzyme B reversed the excess mortality associated with SerpinA3n inhibition, with approximately 92% of mice receiving both ARN1468 and the granzyme B-neutralizing antibody surviving infection **(Fig. 6E)**. These findings identify granzyme B as a major mediator of the increased susceptibility to WNV-induced mortality associated with pharmacological SerpinA3n inhibition and establish that the protective effects of SerpinA3n are mediated, at least in part, through the inhibition of granzyme B.

## Discussion

The regulation of proteolysis represents an underexplored mechanism of tissue protection during CNS infection. Here, we identify SerpinA3n as an endogenous regulator of inflammatory protease activity in the CNS during viral encephalitis. Loss of SerpinA3n production by either astrocytes or neurons exacerbated mortality and neurovascular injury, while supplementation of SerpinA3n in the CNS was broadly protective. Moreover, neutralization of granzyme B reversed the excess mortality associated with pharmacological SerpinA3n inhibition, directly implicating dysregulated protease activity in the pathogenesis of WNV encephalitis (**Fig. 7**). These findings establish the local balance between proteases and their endogenous inhibitors as an important determinant of disease outcome and identify resident CNS cells as active contributors to the maintenance of proteolytic homeostasis during infection.

**Figure 7.**
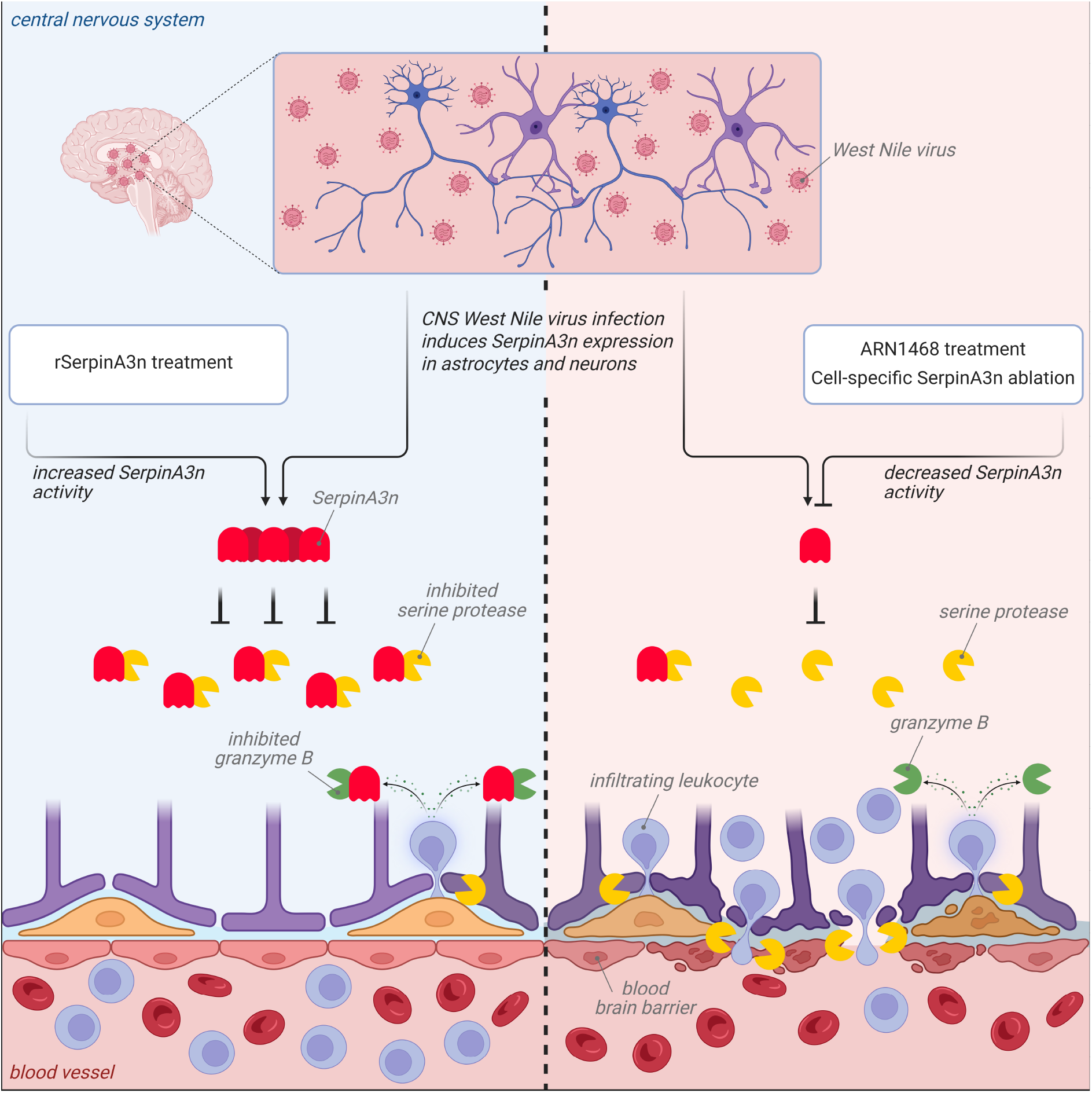
Proposed model of SerpinA3n-mediated protection in WNV encephalitis. Schematic created with BioRender. WNV neuroinvasion induces SerpinA3n expression in astrocytes and neurons, positioning resident CNS cells as local regulators of inflammatory protease activity during infection. *Left*: conditions of increased SerpinA3n activity, modeled by intracerebroventricular (i.c.v.) supplementation with recombinant SerpinA3n (rSerpinA3n). Abundant SerpinA3n forms covalent inhibitory complexes with target serine proteases, including granzyme B released by infiltrating leukocytes at the neurovascular interface. Constraint of extracellular proteolysis preserves blood-brain barrier (BBB) integrity, limits gelatinase/collagenase activity, and restricts further leukocyte accumulation in the CNS. *Right*: conditions of decreased SerpinA3n activity, modeled by i.c.v. treatment with the SerpinA3n inhibitor ARN1468 or by cell-type-specific deletion of SerpinA3n in astrocytes (*Serpina3n^fl/fl^ Aldh1l1*-CreERT2^+^) or neurons (*Serpina3n^fl/fl^ Syn1*-Cre^+^). Insufficient antiprotease activity permits unrestrained serine protease and granzyme B activity, degradation of neurovascular matrix and junctional components, increased BBB permeability, and greater leukocyte infiltration, facilitating a feed-forward cycle of proteolytic injury that culminates in increased mortality.

SerpinA3n appears to protect the infected CNS primarily by limiting immune-mediated tissue injury. Neither supplementation of SerpinA3n nor its deletion from astrocytes or neurons measurably affected CNS viral burden, consistent with our previous observations that neither SerpinA3n nor its upstream regulator, astrocytic RIPK3, directly restricts orthoflavivirus replication [31]. Notably, SerpinA3n supplementation improved survival despite reducing the abundance and relative frequency of WNV-specific CD8⁺ T cells in the CNS, a population with established antiviral functions [54,55]. Although cytotoxic lymphocytes are important for viral clearance, restricting their accumulation or effector activity in the CNS can mitigate neuropathology during orthoflavivirus infection [57,58]. Indeed, CD8⁺ T cell depletion previously rescued the excess mortality associated with SerpinA3n neutralization during ZIKV encephalitis [31]. Local reactivation of infiltrating T cells by antigen-presenting cells is also an important determinant of their effector function during WNV encephalitis [59], and the reduced expression of costimulatory and antigen presentation molecules following SerpinA3n supplementation suggests that SerpinA3n may influence multiple aspects of this response. Collectively, these observations suggest that endogenous antiprotease activity contributes to the balance between protective and pathogenic functions of antiviral immunity in the CNS.

Our identification of granzyme B as a major mediator of neuropathogenesis in the setting of SerpinA3n deficiency is particularly relevant given its established contribution to WNV clearance from the CNS [44–47]. SerpinA3n has previously been shown to protect neurons from granzyme B-mediated injury [60], and granule-derived granzyme B contributes substantially to T cell-mediated killing of human neurons [61]. However, the pathogenic consequences of granzyme B activity may extend well beyond its canonical intracellular cytotoxic functions. Extracellular granzyme B cleaves components of the extracellular matrix, including fibronectin, laminin, and proteoglycans, potentially compromising neurovascular integrity and increasing the susceptibility of the vascular matrix to further proteolytic degradation [14–16]. Granzyme B can also activate protease-activated receptor-1 (PAR1) on neurons, eliciting neurotoxicity through PLCβ–IP3 signaling [17]. Local secretion of SerpinA3n by resident CNS cells may therefore be particularly important for restricting extracellular granzyme B activity, where proteolysis is less tightly coupled to the recognition of infected cells. Defining the relative contributions of extracellular and perforin-dependent intracellular granzyme B activity to viral neuropathogenesis will be an important direction for future studies.

An additional implication of our findings is that serpins may regulate proteolytic activity in the CNS beyond the immediate repertoire of their protease targets. Although SerpinA3n is not known to directly inhibit zinc-dependent MMPs [56], manipulation of SerpinA3n abundance consistently influenced gelatinase/collagenase activity in the infected CNS in our study. MMP-9 is induced during orthoflavivirus encephalitis and contributes to neuroinvasion and BBB disruption [62,63], while focal MMP-2 and MMP-9 activity at the BBB can facilitate leukocyte migration into the CNS [64]. Given that infiltrating neutrophils and monocytes are important sources of MMPs [65,66], SerpinA3n may indirectly regulate MMP activity by limiting the accumulation of these cells. Additionally, neutrophil elastase and cathepsin G, both targets of SerpinA3n [28], can also activate latent MMPs [8,67,68], suggesting that SerpinA3n may act upstream of proteolytic cascades that encompass multiple classes of proteases. This mechanism may help explain the elevated gelatinase/collagenase activity observed in mice lacking astrocytic SerpinA3n in the absence of increased leukocyte accumulation. Although the identities of the proteases responsible for the increased gelatinolytic activity observed in our study remain to be established, these findings highlight the potential for endogenous serpins to exert broad control over proteolysis through the regulation of protease activation cascades.

Our findings also underscore the importance of the cellular and inflammatory contexts in which serpins are expressed. Neuron- and astrocyte-derived SerpinA3n have been ascribed divergent functions in models of CNS injury and disease [30]. Neuronal SerpinA3n protects neurovascular integrity following cerebral ischemia and traumatic brain injury through inhibition of granzyme B and neutrophil elastase, respectively [69,70]. Conversely, astrocytic SerpinA3n has been implicated in the pathogenesis of temporal lobe epilepsy and in the secretory phenotype of senescent astrocytes [71,72], while human SERPINA3 has been implicated in the propagation of inflammatory signaling at the BBB [73]. The biological consequences of serpin induction likely depend in part on the repertoire, abundance, and distribution of active proteases in a given inflammatory environment. Even within the context of WNV encephalitis, neuronal and astrocytic SerpinA3n exhibited distinct immunoregulatory functions despite their shared contributions to host protection. Differences in the anatomical distribution, subcellular localization, or kinetics of SerpinA3n secretion by these cell populations may determine its access to particular pools of inflammatory proteases and their substrates. Understanding how the spatial organization of serpin expression and protease activity influences neuroinflammation may help reconcile the diverse functions attributed to SerpinA3n across CNS pathologies.

Taken together, our findings support a model in which astrocyte- and neuron-derived SerpinA3n restrains inflammatory protease activity at the neurovascular interface, preserving BBB integrity and limiting immune-mediated tissue injury during WNV encephalitis **(Fig. 7)**. More broadly, our findings suggest that restoring proteolytic homeostasis may represent a therapeutic strategy for limiting neuropathology during CNS infection without broadly suppressing antiviral immunity. Both augmentation of endogenous antiprotease activity and selective inhibition of pathogenic effector proteases warrant consideration, particularly given ongoing efforts to develop pharmacological inhibitors of extracellular granzyme B for other inflammatory conditions [74]. Defining the protease networks mobilized during CNS infection, and the endogenous mechanisms that constrain their activity, may reveal new opportunities to limit immune-mediated tissue injury across diverse infectious and inflammatory diseases of the CNS.

## Materials and Methods

### Mice

All mice were housed under specific pathogen-free conditions in Nelson Biological Laboratories at Rutgers University and maintained on a mixed C57BL/6J × 129S genetic background. Adult mice of both sexes (>8 weeks of age) were used in all experiments in accordance with protocols approved by the Rutgers University Institutional Animal Care and Use Committee (protocol 201900016). *Serpina3n*^fl/fl^ (#027511), *Aldh1l1*-CreERT2 (#031008), and *Syn1*-Cre (#003966) mice were obtained from The Jackson Laboratory. The *Syn1*-Cre line expresses Cre recombinase constitutively, whereas Cre recombinase activity in the *Aldh1l1*-CreERT2 line is tamoxifen-inducible. All mice used in experiments involving the *Aldh1l1*-CreERT2 line, including Cre-negative littermate controls, received daily intraperitoneal (i.p.) injections of tamoxifen (60 mg/kg; Sigma-Aldrich, #T5648) in sunflower oil (Sigma-Aldrich, #S5007) for five consecutive days. Experiments were initiated 1–2 weeks after the final tamoxifen administration.

### Viruses and virologic assays

WNV strain WN02-Bird 114 [75] was generously provided by Dr. Bobby Brooke Herrera (Rutgers Robert Wood Johnson Medical School, New Brunswick, NJ, USA). Viral stocks were generated by infecting Vero cells (ATCC, #CCL-81) at a multiplicity of infection (MOI) of 0.01 and harvesting supernatants at 72 hours after infection. Viral titers of stocks were determined by plaque assay on Vero cells. Cells were maintained in DMEM (Corning, #10-013-CV) supplemented with 10% heat-inactivated fetal bovine serum (FBS, Gemini Biosciences #100-106), 1% Penicillin-Streptomycin-Glutamine (Gemini Biosciences #400-110), 1% Amphotericin B (Gemini Biosciences #400-104), 1% Non-Essential Amino Acids (Cytiva, #SH30238.01), and 1% HEPES (Cytiva, #SH30237.01). Plaque assay medium was composed of 1X EMEM (Lonza #12-684F) supplemented with 2% heat-inactivated FBS (Gemini Biosciences, #100-106), 1% Penicillin-Streptomycin-Glutamine (Gemini Biosciences, #400-110), 1% Amphotericin B (Gemini Biosciences #400-104), 1% Non-Essential Amino Acids (Cytiva, #SH30238.01), 1% HEPES (Cytiva, #SH30237.01), 0.75% Sodium Bicarbonate (VWR, #BDH9280), and 0.5% Methyl Cellulose (VWR, #K390). Plaque assays were performed by adding 100 μl of serially diluted sample to 12-well plates containing 200,000 Vero cells per well and incubating for 1 hour at 37 °C. Plates were incubated with plaque assay medium at 37°C and 5% CO2 for 3 days. After incubation, the overlay medium was removed, and the plates were stained/fixed with 10% neutral-buffered formalin (VWR, #89370) and 0.25% crystal violet (VWR, #0528) for 20 to 30 min. Plates were washed repeatedly in water and allowed to dry before counting visible plaques.

### Mouse infections and tissue harvesting

Isoflurane anesthesia was used for all procedures. Mice were inoculated subcutaneously in a rear footpad (50 μl) with 1875 PFU of WNV-WN02-Bird114. Infected mice were monitored daily by a blinded operator for weight loss and clinical signs of disease, including hunched posture, ruffled fur, hindlimb weakness, and paresis. Mice reaching a moribund state or losing more than 20% of their initial body weight were euthanized, and the date of euthanasia was recorded for the purpose of survival studies. For brain harvest experiments, mice underwent cardiac perfusion with 30 mL of cold sterile 1X phosphate-buffered saline (PBS). For regional analyses, the cortex, cerebellum, brainstem, diencephalon, and spinal cord were dissected in some experiments, as indicated in the text. All tissues were homogenized with 1.0-mm zirconia/silica beads (BioSpec Products, #11079110z) in a Bead Ruptor Elite (Omni International).

### Quantitative real-time PCR

Total RNA was isolated from tissue homogenates using the Direct-zol RNA Miniprep Kit (Zymo Research, #R2051) according to the manufacturers’ instructions. RNA concentration and purity were assessed using a SpectraMax QuickDrop spectrophotometer (Molecular Devices). Within each experiment, RNA concentrations were normalized across samples prior to reverse transcription using the qScript cDNA Synthesis Kit (Quantabio, #95047). Quantitative real-time PCR (qRT-PCR) was performed using SYBR Green Master Mix (Bio-Rad, #1725275) on a QuantStudio 5 instrument (Applied Biosystems). Cycle threshold (Ct) values were normalized to those of 18S rRNA (ΔCt = Ct_target − Ct_18S). Primer sequences are provided in **Table S1**.

### Flow cytometry

Whole brains were harvested from perfused mice and collected in cold Hanks’ balanced salt solution (HBSS; VWR, #02-1231-0500) supplemented with 10 mM HEPES (Cytiva, #SH30237.01). Brain tissue was enzymatically dissociated using collagenase type I (Sigma-Aldrich, #C0130) and deoxyribonuclease I (Sigma-Aldrich, #D4527), followed by mechanical dissociation and filtration through a 70-µm cell strainer. Leukocytes were isolated by Percoll density gradient centrifugation (Cytiva, #17-0891-02). Following red blood cell (RBC) lysis (Tonbo Biosciences, #TNB4300-L100), cells were washed and resuspended in fluorescence-activated cell sorting (FACS) buffer consisting of 1× PBS supplemented with 0.2% sodium azide, 5% FBS, 2 mM EDTA, and 10 mM HEPES.

Leukocytes were stained with Zombie NIR viability dye (BioLegend, #423105) for 20 min at 4°C, followed by incubation with 0.5% normal mouse serum and 1% FcX Block (BioLegend, #101320) in FACS buffer for 15 min at 4°C. Cells were subsequently stained with fluorophore-conjugated antibodies against CD45.2 (BioLegend, clone 104, #109821), MHC-II (I-A/I-E; BioLegend, clone M5/114.15.2, #107641), CD11b (BioLegend, clone M1/70, #101224), CD3 (BioLegend, clone 17A2, #100259), CD4 (BioLegend, clone RM4-5, #100557), CD8α (BioLegend, clone 53-6.7, #100752), CD19 (Invitrogen, clone eBio1D3, #M004T03B12-A), CD11c (BioLegend, clone N418, #117326), F4/80 (BioLegend, clone BM8, #123145), Ly-6C (BioLegend, clone HK1.4, #128030), Ly-6G (BioLegend, clone 1A8, #127645), CD80 (BioLegend, clone 16-10A1, #104759), and NKp46 (Invitrogen, clone 29A1.4, #50-3351-82). For identification of WNV-specific CD8⁺ T cells, H-2Dᵇ tetramers loaded with the immunodominant WNV NS4B peptide (SSVWNATTA; residues 2488–2496 of the viral polyprotein), provided by the NIH Tetramer Core Facility (Emory University, Atlanta, GA), were included in the surface antibody staining panel. Cells were stained for 20 min at 4°C, washed in FACS buffer, and fixed in 1% paraformaldehyde (PFA) in PBS.

Data were acquired using a Cytek Aurora spectral flow cytometer (Cytek Biosciences) and analyzed using FlowJo software (Tree Star). Absolute cell numbers were determined using counting beads (Thermo Fisher Scientific, #C36995). Splenocytes were prepared by mechanical dissociation of spleens through a 70-µm cell strainer, followed by RBC lysis and washing in FACS buffer. Splenocytes and compensation beads (Thermo Fisher Scientific, #01-2222-42) were used to generate single-color controls.

### In vivo assessment of BBB permeability

BBB permeability was assessed by quantifying sodium fluorescein extravasation into CNS tissues. Mice received i.p. injections of fluorescein sodium salt (100 µL of a 100 mg/mL solution in sterile PBS; Sigma-Aldrich, #F6377). After 45 min, blood was collected, and mice were perfused with PBS. Serum and clarified CNS tissue homogenates were mixed 1:1 with 2% trichloroacetic acid (Sigma-Aldrich, #T0699) and incubated overnight at 4°C. Precipitated proteins were removed by centrifugation at 2,823 × g for 10 min at 4°C. Supernatants were mixed 1:1 with borate buffer (pH 11; Sigma-Aldrich, #1094621000) to neutralize the pH. Fluorescence was measured in black-walled, optically clear 96-well plates (Corning, #3904) using a SpectraMax i3x multimode plate reader (Molecular Devices). Tissue fluorescence values were normalized to the corresponding serum fluorescence values for each mouse.

### In vivo drug treatments

Systemic ARN1468 treatment: Mice were injected i.p. with ARN1468 (2.7 mg/kg; MedChemExpress, #HY-150514) dissolved in 10% PEG400, 10% Tween 80, and 80% saline (v/v). Injections were given twice daily (morning and evening) from 4 to 8 dpi. Control mice received vehicle on the same schedule.

Intracerebroventricular injections: Under isoflurane anesthesia, mice received i.c.v. injections (−0.3 anterior-posterior; +1.0 medio-lateral; +2.5 dorso-ventral). Mice received one of the following, diluted in artificial cerebrospinal fluid (ACSF; Tocris Bioscience, #3525): (i) 5 µg ARN1468 in 10% DMSO/90% ACSF; (ii) 4 ng recombinant mouse SerpinA3N (rSerpinA3N; R&D Systems, #4709-PI); (iii) 2 µg goat anti-mouse Granzyme B antibody (R&D Systems, #AF1865); or (iv) 2 µg goat IgG isotype control (R&D Systems, #AB-108-C).

### SerpinA3n ELISA

SerpinA3n abundance in CNS tissue homogenates was assessed using a mouse SerpinA3n ELISA kit (RayBiotech, #ELM-SERPINA3N-1) according to the manufacturer’s instructions. Absorbance at 450 nm was measured using a SpectraMax i3x plate reader (Molecular Devices). Absorbance values within the linear range were normalized to those of uninfected (0 dpi) controls.

### Gelatinase/collagenase activity assay

Gelatinase/collagenase activity in brain homogenates was measured using the EnzChek Gelatinase/Collagenase Assay Kit (Thermo Fisher Scientific, #E12055) according to the manufacturer’s instructions. Brain homogenates were centrifuged at 15,000 × g for 10 min at 4°C, and clarified supernatants were incubated with a fluorogenic gelatin substrate in black-walled 96-well plates for 2 h at room temperature. Fluorescence was measured using a SpectraMax i3x plate reader (Molecular Devices) at excitation and emission wavelengths of 490 and 515 nm, respectively.

### Granzyme B activity assay

Granzyme B activity in brain homogenates was measured using the QuickZyme Granzyme B Activity Assay Reagent Set (QuickZyme Biosciences, #QZBGRB2) according to the manufacturer’s instructions. Brain homogenates were centrifuged at 15,000 × g for 10 min at 4°C, and clarified supernatants were transferred to antibody-coated plates to capture granzyme B. Enzymatic activity was assessed by cleavage of a chromogenic substrate over 24 h at 37°C. Absorbance was measured at 405 nm using a SpectraMax i3x plate reader (Molecular Devices).

### Statistical analysis

Statistical analyses were performed using GraphPad Prism 11 (GraphPad Software). Survival curves were compared using the log-rank (Mantel–Cox) test. Other comparisons were performed using two-tailed Student’s *t* tests or one-or two-way analysis of variance (ANOVA) followed by Dunnett’s, Šídák’s, or Fisher’s least significant difference (LSD) multiple-comparison tests, as indicated in the figure legends. Correlations were assessed using Spearman’s rank correlation test. Differences were considered statistically significant at *P* < 0.05. Unless otherwise indicated, individual data points represent biological replicates obtained from distinct mice, data are presented as mean ± SEM, and data were collected from at least two independent experiments. No data points were excluded from statistical analyses.

## Acknowledgments

We thank the NIH Tetramer Core Facility for providing WNV NS4B-specific H-2Dᵇ tetramers.

## Author contributions (CRediT taxonomy)

Conceptualization: VK and BPD; Investigation: VK, EM, JPA, ED, CA, and BPD; Formal analysis: VK and BPD; Visualization: VK, EM, and BPD; Writing – original draft: VK; Writing – review & editing: VK and BPD; Supervision: CA and BPD; Funding acquisition: BPD.

---

**Figure S1.**
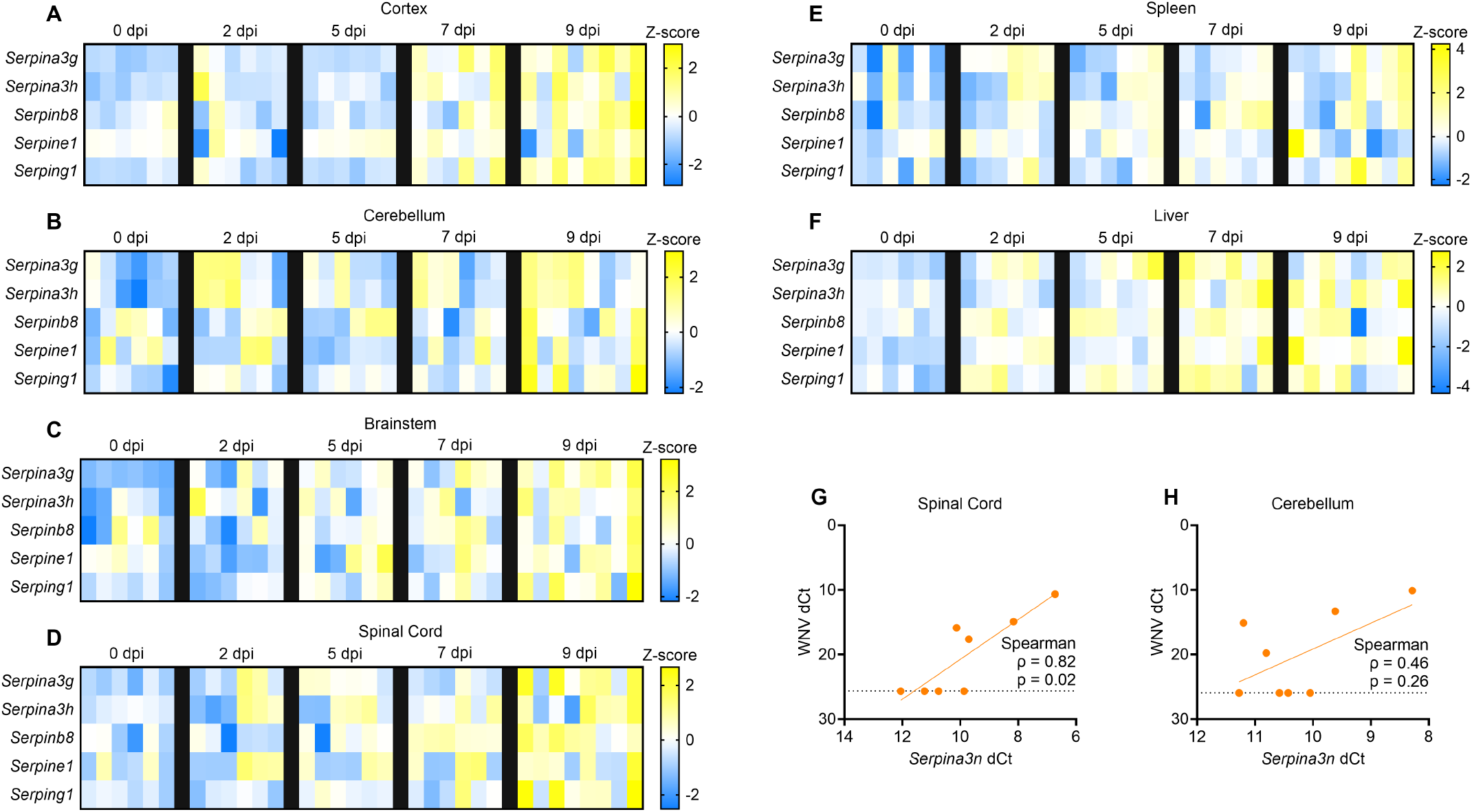
Expression of serpin family members and correlation of *Serpina3n* expression with WNV RNA in the cerebellum and spinal cord during WNV infection. (A–F) Heatmaps depicting *Serpina3g*, *Serpina3h*, *Serpinb8*, *Serpine1*, and *Serping1* transcript levels in the cortex (A), cerebellum (B), brainstem (C), spinal cord (D), spleen (E), and liver (F) at indicated days post-infection (dpi). Z-scores were calculated from ddCt values. Each square represents an individual mouse, and each row represents a gene. (G, H) Correlation between *Serpina3n* dCt and WNV dCt in individual mice at 9 dpi in the spinal cord (G) and cerebellum (H). Spearman correlation coefficients (ρ) and p-values are shown. The linear regression line is shown for visualization only. The dotted line marks the limit of detection for WNV RNA; data points on this line are below the limit of detection. n = 6 mice per group at 0, 2, 5, and 7 dpi and n = 8 mice at 9 dpi.

**Figure S2.**
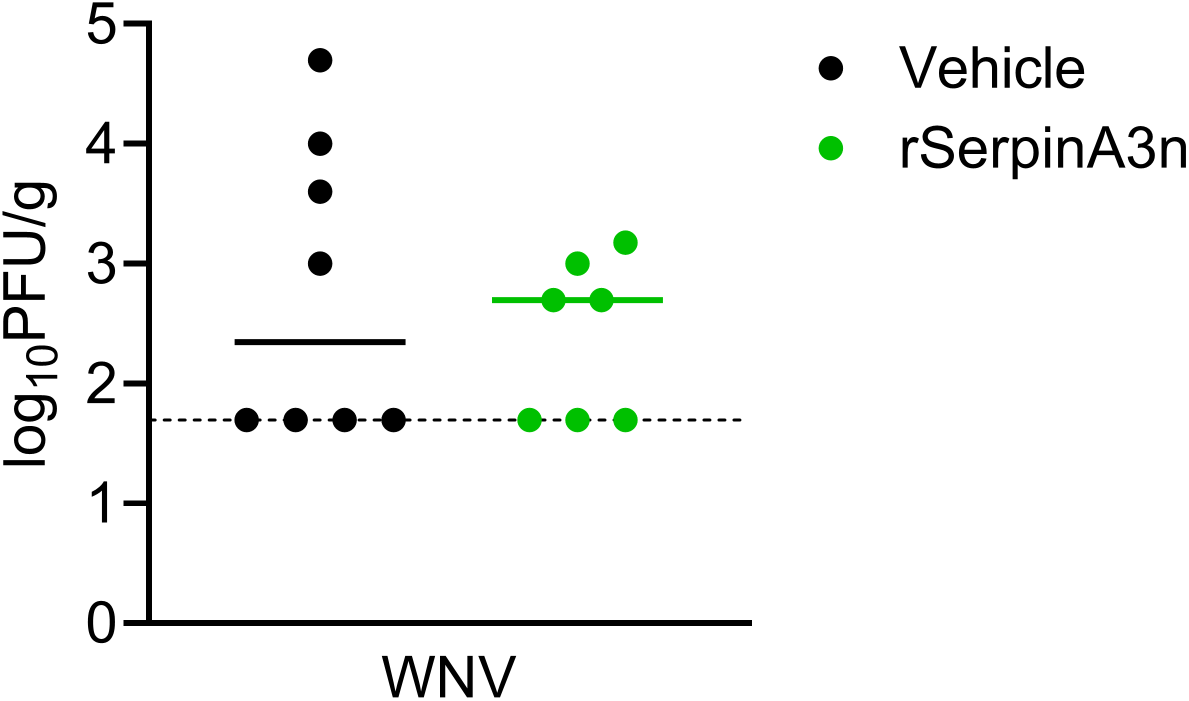
CNS-localized rSerpinA3n treatment does not alter brain viral burden in WNV-infected mice. Infectious viral titers in the brain at 8 days post-infection (dpi), measured by plaque assay, in WNV-infected mice treated intracerebroventricularly with vehicle or recombinant SerpinA3n (rSerpinA3n) at 5 and 7 dpi. Titers are expressed as log_10_PFU per gram of tissue. Each dot represents an individual mouse, and horizontal lines indicate the median. The dotted line marks the limit of detection of the plaque assay; data points on this line are below the limit of detection. Groups were compared by Welch’s t test. n = 8 for Vehicle and n = 7 for rSerpinA3n.

**Figure S3.**
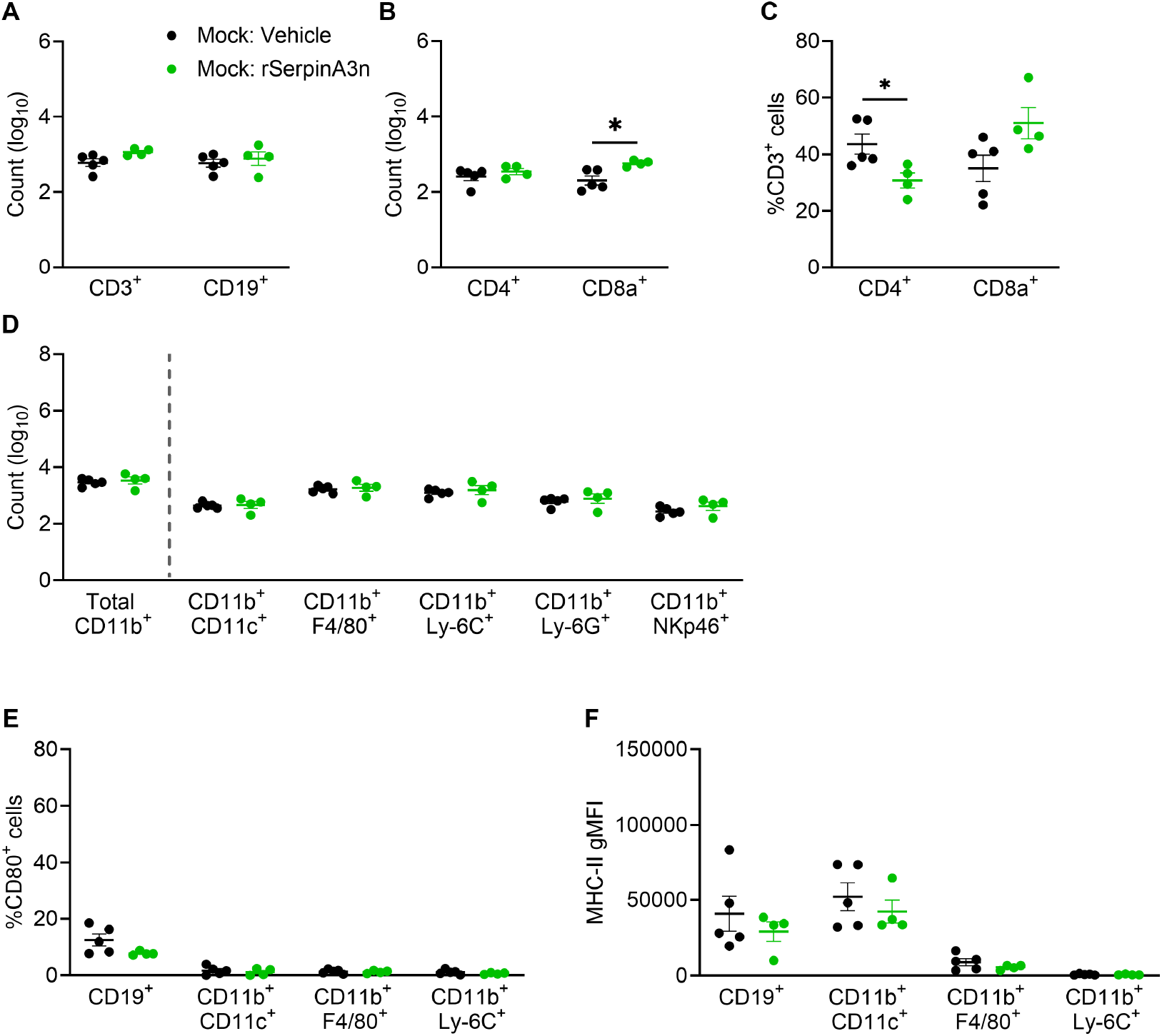
rSerpinA3n has minimal effect on CNS leukocyte recruitment and antigen-presenting cell activation in mock-infected mice. Flow cytometry analysis of brain leukocytes from mock-infected mice treated intracerebroventricularly with vehicle or recombinant SerpinA3n (rSerpinA3n), analyzed with the same gating strategy as in Figure 3. (A) Numbers of CD3^+^ T cells and CD19^+^ B cells, gated on CD45.2^+^ cells. (B, C) Numbers (B) and percentages among CD3^+^ cells (C) of CD4^+^ and CD8^+^ T cells, gated on CD3^+^ cells. (D) Numbers of total CD11b^+^ cells (gated on CD45.2^+^ cells) and of the indicated innate populations: CD11b^+^CD11c^+^, CD11b^+^F4/80^+^, CD11b^+^Ly-6C^+^, CD11b^+^Ly-6G^+^, and CD11b^+^NKp46^+^ cells (each gated on CD11b^+^ cells). (E) Percentage of CD80^+^ cells among CD19^+^ B cells (gated on CD45.2^+^ cells) and among CD11b^+^CD11c^+^, CD11b^+^F4/80^+^, and CD11b^+^Ly-6C^+^ cells (each gated on CD11b^+^ cells). (F) MHC-II geometric mean fluorescence intensity (gMFI) in the same populations as in E. In A, B, and D, cell counts are shown on a log_10_ scale. Each dot represents an individual mouse. Data are presented as mean ± SEM. Vehicle and rSerpinA3n groups were compared within each cell population by Student’s unpaired t tests. *p < 0.05. n = 5 for Mock:Vehicle and n = 4 for Mock:rSerpinA3n.

**Figure S4.**
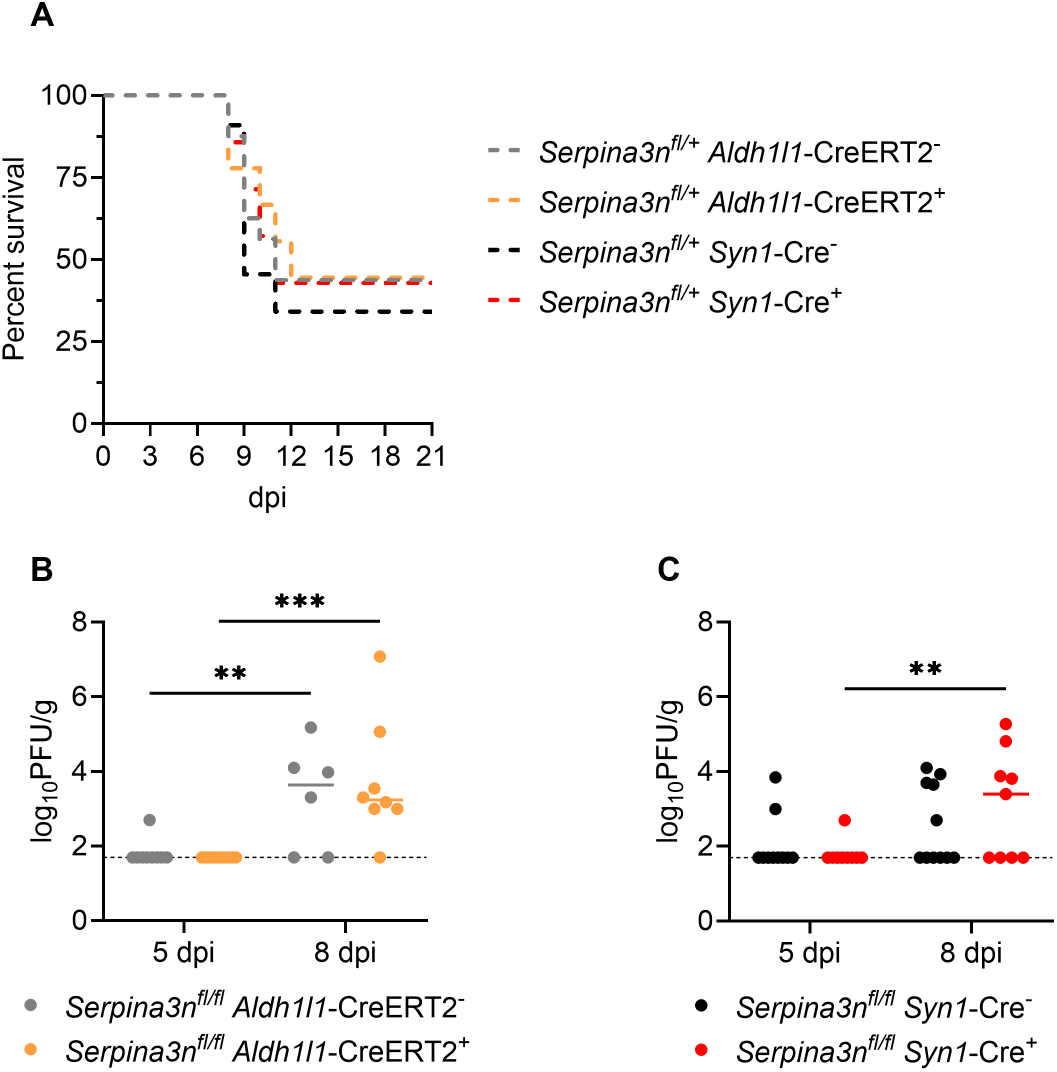
Heterozygous deletion of astrocytic or neuronal SerpinA3n does not alter survival, and homozygous deletion does not change CNS viral burden during WNV infection. (A) Survival of WNV-infected *Serpina3n^fl/+^ Aldh1l1*-CreERT2 and *Serpina3n^fl/+^ Syn1*-Cre mice (Cre-positive) and their Cre-negative littermate controls. Survival was compared by log-rank (Mantel-Cox) test, with Cre-positive mice compared only with Cre-negative mice of the same Cre line. (B, C) Infectious viral titers in the CNS at 5 and 8 days post-infection (dpi), measured by plaque assay and expressed as log_10_ PFU per gram of tissue, in *Serpina3n^fl/fl^ Aldh1l1*-CreERT2 (B) and *Serpina3n^fl/fl^ Syn1*-Cre (C) mice and their Cre-negative littermate controls. Each dot represents an individual mouse, and horizontal lines indicate the median. The dotted line marks the limit of detection of the plaque assay; data points on this line are below the limit of detection. A two-way ANOVA was performed with the uncorrected Fisher’s LSD test for the following comparisons: Cre-negative vs. Cre-positive at 5 dpi and at 8 dpi, and 5 dpi vs. 8 dpi within each genotype. Only significant comparisons are shown. **p < 0.01, ***p < 0.001. Sample sizes: (A) *Serpina3n^fl/+^ Aldh1l1*-CreERT2: Cre-negative n = 16, Cre-positive n = 9; *Serpina3n^fl/+^ Syn1*-Cre: Cre-negative n = 11, Cre-positive n = 7. (B) *Serpina3n^fl/fl^ Aldh1l1*-CreERT2: 5 dpi, Cre-negative n = 9 and Cre-positive n = 9; 8 dpi, Cre-negative n = 6 and Cre-positive n = 8. (C) *Serpina3n^fl/fl^ Syn1*-Cre: 5 dpi, Cre-negative n = 10 and Cre-positive n = 9; 8 dpi, Cre-negative n = 11 and Cre-positive n = 9.

**Table S1.**
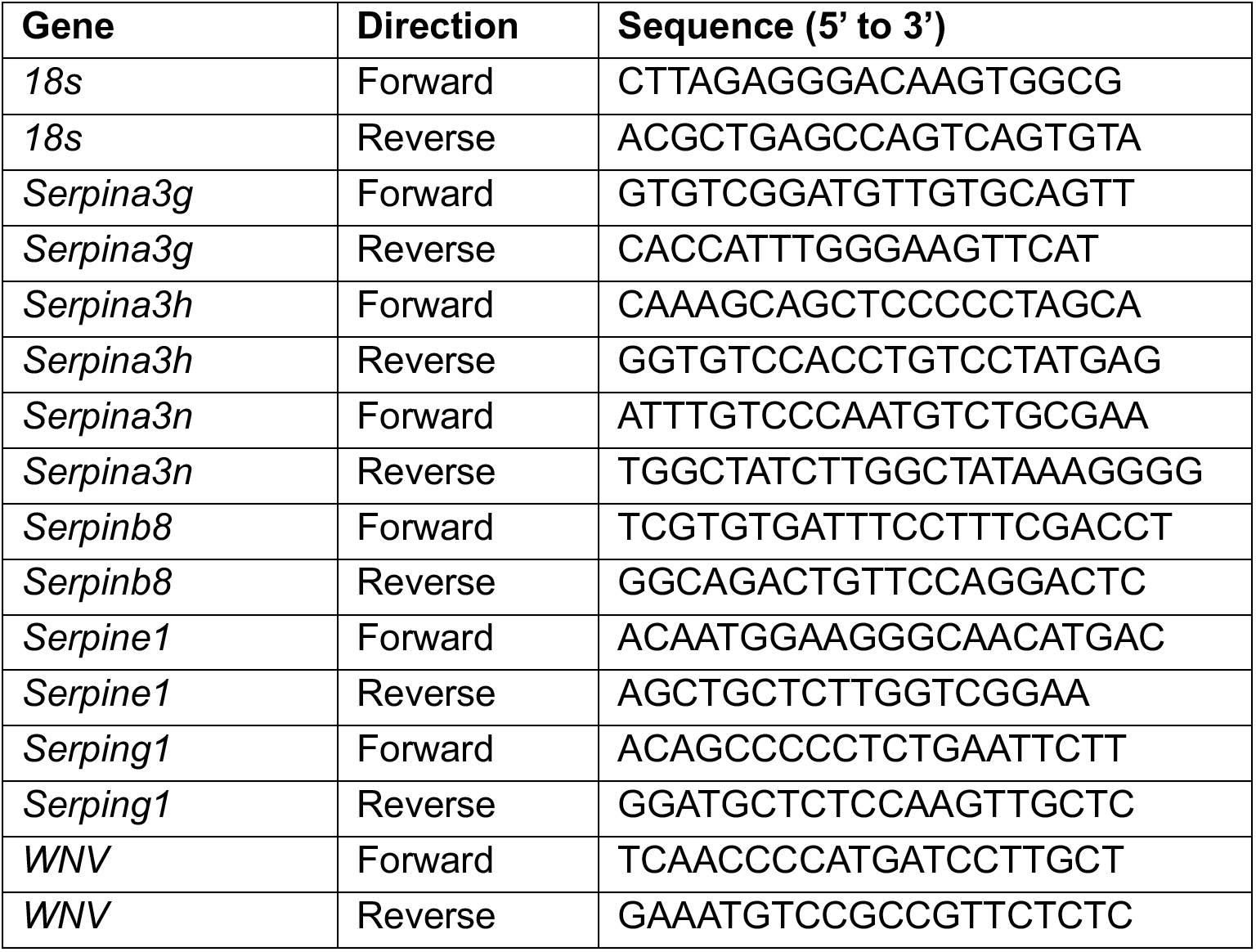
Primer sequences for qRT-PCR analysis.

